# Parkinson’s disease-derived α-Synuclein assemblies combined with chronic-type inflammatory cues promote a neurotoxic microglial phenotype

**DOI:** 10.1101/2023.11.30.569208

**Authors:** Cansu Yildirim, Alexis Fenyi, Pierre Besnault, Lina Gomez, Julia E. Sepulveda-Diaz, Patrick P. Michel, Ronald Melki, Stéphane Hunot

## Abstract

Parkinson’s disease (PD) is a common age-related neurodegenerative disorder characterized by the aggregation of α-synuclein (αSYN) building up intraneuronal inclusions termed Lewy pathology. Mounting evidence suggests that neuron-released αSYN aggregates could be central to microglial activation, which in turn mounts and orchestrates neuroinflammatory processes potentially harmful to neurons. Therefore, understanding the mechanisms that drive microglial cell activation, polarization and function in PD might have important therapeutic implications. Here, using primary microglia, we investigated the inflammatory potential of pure αSYN fibrils derived from PD patients. We further explored and characterized microglial cell responses to a chronic-type inflammatory stimulation combining PD patient-derived αSYN fibrils (F^PD^), Tumor necrosis factor-α (TNFα) and prostaglandin E_2_ (PGE_2_) (TPF^PD^). We showed that F^PD^ hold stronger inflammatory potency than pure αSYN fibrils generated *de novo*. When combined with TNFα and PGE_2_, F^PD^ polarizes microglia toward a particular functional phenotype departing from F^PD^-treated cells and featuring lower inflammatory cytokine and higher glutamate release. Whereas metabolomic studies showed that TPF^PD^-exposed microglia were closely related to classically activated M1 proinflammatory cells, notably with similar tricarboxylic acid cycle disruption, transcriptomic analysis revealed that TPF^PD^-activated microglia assume a unique molecular signature highlighting upregulation of genes involved in glutathione and iron metabolisms. In particular, TPF^PD^-specific upregulation of *Slc7a11* (which encodes the cystine-glutamate antiporter xCT) was consistent with the increased glutamate response and cytotoxic activity of these cells toward midbrain dopaminergic neurons *in vitro*. Together, these data further extend the structure-pathological relationship of αSYN fibrillar polymorphs to their innate immune properties and demonstrate that PD-derived αSYN fibrils, TNFα and PGE_2_ act in concert to drive microglial cell activation toward a specific and highly neurotoxic chronic-type inflammatory phenotype characterized by robust glutamate release and iron retention.

## Introduction

Parkinson’s disease (PD) is a common and multisystem neurodegenerative disorder characterized by both motor and nonmotor symptoms. The neuropathological hallmark of PD is the progressive and massive degeneration of dopaminergic neurons (DNs) in the *substantia nigra* associated with α-Synuclein (αSYN)-rich deposits building up cytoplasmic inclusions known as Lewy pathology (LP). The aggregation of αSYN may promote several molecular and biochemical defects affecting key cellular functions and driving the progressive demise of DNs [1]. In addition to these toxic cell-autonomous mechanisms, pathological αSYN assemblies can also spread from cell to cell to seed aggregate formation in newly contaminated neurons. This prion-like mechanism is believed to contribute to the progression of αSYN pathology across brain networks over time and to be linked to the progressive clinical deterioration of patients [2, 3]. Accordingly, released extracellular αSYN aggregates may directly interact with and activate surrounding glial cells including microglia to initiate a deleterious inflammatory response. Indeed, evidence from epidemiological, *postmortem*, and animal studies suggests that neuroinflammatory processes associated with glial and immune cell activation participate in the progression of DN cell death [4].

Microglia, the most abundant tissue macrophages in the brain, execute numerous physiological functions important for the maintenance of tissue homeostasis, synapse remodeling, neurotrophic factor secretion and clearance of debris. However, under neurodegenerative conditions, and more particularly upon chronic exposure to aberrant proteins, microglial cells mount persistent sterile (as opposed to infectious) and proinflammatory immune responses [5]. In PD, major progress in our understanding of microglial cell activation and related inflammation has shown that αSYN assemblies behave as true molecular motifs associated with microbes (pathogen-associated molecular patterns or PAMPs). These motifs are recognized by innate immune cells thanks to a panel of immune receptors recognizing these molecular patterns (pattern recognition receptors or PRRs) such as Toll-like receptors (TLRs). Thus, mounting evidence suggests that neuron-released αSYN assemblies could be central to microglial cell activation and pathological inflammatory responses through the activation of TLRs and other immune-related receptors including CD36 [6-9]. However, these studies also revealed that the set of receptors involved in this activation crucially depends on the type of αSYN assemblies used in the different experimental setups. Indeed, αSYN can assemble into structurally distinct fibrillar assemblies (also termed polymorphs or strains) that exhibit different phenotypic traits [10-12]. As we previously showed, these strains exhibit distinct spreading, tropism, clearance, and neurotoxicity in cells and in the rodent brain, which might explain the pathological and phenotypic diversity of different synucleinopathies such as PD, multiple system atrophy and dementia with Lewy bodies [10-14]. Importantly, we recently strengthened this concept using patient-derived αSYN strains generated by protein misfolding cyclic amplification (PMCA) [15]. Besides demonstrating a structure-pathology relationship we brought evidence for differences between patient-derived and *de novo*-generated αSYN strains [15]. Taken together, these data suggest that the intrinsic structure of fibrillar αSYN is a major determinant of its biological properties. A likely explanation for this structure-function relationship is that differences in surface accessibility contribute to the differential binding of distinct αSYN strains to partner proteins such as cell surface receptors [12, 16]. Hence, whether patient-derived αSYN strains exhibit inflammatory activity similar to or different from that of previously described *de novo*-generated αSYN polymorphs is unknown, but their use in model systems has the potential to better model disease-associated microglial activation and perhaps contribute to more relevant knowledge.

An often neglected, but important issue for understanding the pathomechanisms linked to innate inflammation in PD is to comprehensively define the complex network of stimuli that drive microglial cell activation and function. Most cell culture models for studying microglia-associated immunity in PD have been developed upon exposure of microglia to the prototypical bacterial inflammogen lipopolysaccharide (LPS) or, much elegantly, to different types of recombinant αSYN aggregates of various purities as disease-specific triggers and TLR-activating pathological ligands [8, 9, 17, 18]. Although these models have brought invaluable knowledge about the molecular mechanisms and pathways involved in microglial cell activation, they are limited in that they do not consider specific aspects of PD-related neuroinflammation which is chronic in nature and most likely involves the combined action of several mediators in the inflammatory milieu. It is now well established that the phenotypes and functions that activated microglia can adopt are diverse and strongly depend on the nature, intensity and complexity of stimulation in response to various cues [19]. Far from the original idea that macrophages can be polarized into classically (M1) or alternatively (M2) activated cells representing two polar extremes of molecular programming, there is now compelling evidence suggesting that they can adopt a much larger spectrum of activation states associated with different phenotypes and, presumably, functions [20, 21]. Notably, it has been shown that human macrophages engage in a specific molecular program and adopt a unique phenotype when exposed to a chronic-type inflammatory stimulus composed of Tumor necrosis factor (TNF)-α, prostaglandin E_2_ (PGE_2_) and TLR2 ligands such as the synthetic molecule Pam3CSK4 (referred to as “TPP” stimulation) [21]. These inflammatory factors are commonly found in chronic inflammatory conditions such as *Mycobacterium tuberculosis*-associated granulomatous diseases or granulomatous listeriosis [22-26]. Remarkably, the expression of these factors is elevated in the brains of PD patients as well; αSYN aggregates being the most likely pathological ligand of TLR2 whose expression is tightly linked to PD pathogenesis [27-29], suggesting that they may act in concert to shape microglial activation into a unique disease phenotype. Yet, cell-specific immune properties might be important and whether microglial cells behave similarly to peripheral macrophages upon chronic-type inflammatory stimulation has not been explored.

In this study, using a model system of cultured primary microglial cells, we sought to document the inflammatory properties of patient-derived αSYN strains and to determine whether microglia exposed to PD-relevant inflammatory cues in a novel and original combinatory approach adopt a specific molecular phenotype and function. We found that PD patient-derived αSYN fibrils hold stronger inflammatory properties than αSYN fibrils assembled *de novo*. We also showed that, compared to classical M1-shaped microglia (LPS-induced), cells simultaneously exposed to PD patient-derived fibrils, TNFα and PGE_2_ assume a unique chronic-type inflammatory molecular signature associated with increased glutamate release and iron retention propensities as well as higher neurotoxic activity.

## Methods

### Animals

Animal care and housing were conducted in accordance with the recommendations of the European Union Council Directives (2010/63/EU). C57BL/6J mice were obtained from Janvier LABS (Le Genest St Isle, France).

### Reagents and antibodies

Leibovitz’s L-15 medium (Thermo Fisher Scientific, Illkirch, France, #11415049), Dulbecco’s Modified Eagle Medium/F-12 nutrient mixture (DMEM; Thermo Fisher Scientific, #31885023), Neurobasal-A medium (Nb; Thermo Fisher Scientific, #10888022), B27 supplement minus antioxidants (Thermo Fisher Scientific, #10889038), N2 mix (Thermo Fisher Scientific, #17502048), RPMI-1640 Glutamax (Invitrogen, #61870010), RPMI-1640 (Thermo Fisher Scientific, #21875034), penicillin/streptomycin (Thermo Fisher Scientific, #15140122), antibiotic/antimycotic solution (Invitrogen, #15240062), Trypsin 0.05% EDTA (Thermo Fisher Scientific, #25300054), polyethylenimine (PEI; Sigma Aldrich, #P3143), Histopaque-1077 (Sigma Aldrich, #10771-500M), fetal bovine serum (FBS; Biowest, #S1820-500), Nitrotetrazolium Blue chloride (Sigma Aldrich, #N6876), Pam3CSK4 (Invivogen, tlrl-pms), Lipopolysaccharides (LPS) from *Escherichia coli* O26:B6 (Sigma-Aldrich, #L8274), recombinant human TNFα (Sigma Aldrich, #H8916), PGE_2_ (Sigma Aldrich, #P6532), recombinant mouse IFNγ (Immunotools, #12343537), recombinant mouse IL4 (Immunotools, #12340045), recombinant human GM-CSF (R&D, #215-GM-010), recombinant human IL34 (R&D, #5265-IL-010), Sulfasalazine (Sigma Aldrich, S0883), MK-801 (Tocris, #0924), Cytotoxicity Detection Kit (LDH; Roche, #11644793001), Amplex Red Glutamic Acid/Glutamate Oxidase Kit (Thermo Fisher Scientific, #A12221), NucleoSpin RNA XS, RNA purification kit (Macherey-Nagel, #740902.50), SYBR Green reactifs (LightCycler® 480 SYBR Green I Master; Roche, #04707516001), StraightFrom® LRSC CD14 MicroBead Kit (Miltenyi, #130-117-026), MACS BSA Stock Solution (Miltenyi, #130[7]091[7]376), autoMACS® Rinsing Solution (Miltenyi, #130[7]091[7]222). The following antibodies were used: anti-mouse CD25 (Bio-Rad, MCA1260), anti-mouse CD11b (Bio-Rad, MCA74G), anti-human CCR2 (R&D Systems; MAB150-100), anti-human CX3CR1 (Bio-Rad, AHP566). Mouse TNFα ELISA Kit (Thermo Fisher Scientific, BMS607-3), Mouse IL6 ELISA Kit (Thermo Fisher Scientific, BMS603-2TWO), Mouse IL1β ELISA Kit (Thermo Fisher Scientific, BMS6002), Mouse IL10 ELISA Kit (R&D Systems, M1000B), U-Plex (Meso Scale Discovery, K15069L-1), Mouse Pro-Inflammatory 7-Plex Tissue (Meso Scale Discovery, K15012B-1), Mouse IL1α ELISA Kit (Biolegend, #433404), Mouse CXCL5 ELISA Kit (R&D Systems, MX000), and Hoechst 33342 (Tocris, #5117).

### Human brain tissue collection and preparation of **α**SYN assemblies

Brain tissues from patients suffering from PD (n=4) or DLB (n=4) were obtained at autopsy from the UK Brain Bank (Imperial College London, UK). The clinicopathological description of the patients is detailed elsewhere [15]. After histological identification of LP-rich regions, the cingulate cortex was isolated from the brains and processed into brain homogenates. Patient-derived αSYN assemblies were obtained by protein misfolding cyclic amplification (PMCA) as described elsewhere [15]. Briefly, frozen brain tissues were diluted 1:5 (w/v) in PMCA buffer (150 mM KCl, 50 mM Tris–HCl pH 7.5) and sonicated for 1 min, with 10 s pulses followed by 10 s pauses (SFX 150 Cell Disruptor sonicator equipped with a 3.17 mm Branson microtip probe). Brain homogenates were further diluted in PMCA buffer containing monomeric αSYN (100 µM) to a final concentration of 2% (w:v). PMCA amplification (4 cycles) was performed using a Q700 generator and a 431MPX horn (Qsonica, Newtown, CT, USA) with the following sequence: 15 s of sonication and 5 min pause at 31°C. Cycles 2, 3 and 4 were performed using 1% of the preceding cycle reaction as seeds. At cycle 4, the PMCA reaction products were spun for 30 min at 50,000xg and the pelleted assemblies were resuspended in phosphate-buffered saline (PBS) at a final concentration of 100 µM. Prior to usage, the assemblies were fragmented by sonication for 20 min in 2-ml Eppendorf tubes in a Vial Tweeter powered by an ultrasonic processor UIS250v (250 W, 2.4 kHz; Hielscher Ultrasonic, Teltow, Germany), aliquoted, flash frozen in liquid nitrogen and stored at -80°C. Recombinant αSYN fibrils were prepared and characterized as previously described (Bousset et al., 2013). We used the LALChromogenic Endotoxin Quantitation Kit (Thermo Fisher catalog no. 88282) following manufacturer instructions and a CLARIOstar Plus (BMG Labtech, Ortenberg, Germany) plate reader as described previously [13]. To ascertain that the endotoxin levels were less than 0.015 endotoxin units/mg (EU/mg). In preliminary studies, we observed similar inflammatory activities in cultured microglia from patient-derived αSYN fibrils among patients in each disease group. Therefore, we prepared a single batch of PD and DLB patient-derived αSYN fibrils by mixing equal amount of assemblies generated from the 4 patients in each disease group.

### Transmission electron microscopy

The morphology of the PMCA-amplified αSYN assemblies (3 cycle) was assessed by transmission electron microscopy (TEM) using a Jeol 1400 transmission electron microscope following adsorption onto carbon-coated 200 mesh grids and negative staining with 1% uranyl acetate. The images were acquired using a Gatan Orius CCD camera (Gatan, Elancourt, France).

### Primary mouse microglial culture

Primary mouse microglia were isolated and cultured following a protocol previously described [30]. Newborn pups were sacrificed, and the whole brain was rapidly dissected. After mechanical dissociation of the brain tissue, the cells in suspension were plated onto polyethyleneimine-coated T-75 flasks with 12 ml of DMEM supplemented with 10% FBS and antibiotics (2 brains per flask). At DIV2, the medium was completely changed to fresh medium, and the culture was subsequently left to grow without changing the medium until microglial cell isolation was complete (DIV14–18). For microglial-conditioned medium (MCM) transfer into neuronal culture media, microglia were grown and stimulated in DMEM-based astrocyte-conditioned medium (ACM) prepared as described below.

### Microglial cell seeding and stimulation

Cultured microglia were washed 3 times with DMEM and incubated for 5 min with 5 mL of prewarmed (+37°C) trypsin. DMEM containing 10% FBS (10 mL) was then added to abrogate trypsin activity, and the cells were collected in an ice-cold 50 mL falcon tube before centrifugation at 188 x g for 6 min at 4°C. The cell pellet was then resuspended in DMEM-1% FBS. The cells were plated at a density of 90,000 to 110,000 cells/cm and incubated for 24 h at 37°C for complete settlement. Microglia were stimulated for 24 to 48 hours with LPS (10 ng/mL), Pam3CSK4 (1 µg/mL), TNFα (800 IU/mL), PGE_2_ (1 µg/mL) or fibrillar αSYN assemblies at various concentrations. When indicated, stimulation factors were used in combination.

### Astrocyte-conditioned medium

Astrocyte-conditioned medium (ACM) was prepared from primary cultures of astrocytes obtained from C57BL/6J mouse neonate (P0) brains as described elsewhere [31]. Briefly, brains were dissected and dissociated by mechanical trituration in L15 medium. After centrifugation, the cell pellet was resuspended in DMEM/F-12 plus 10% FBS, and the dissociated cells were distributed and allowed to grow on laminin-coated T-75 flasks. At DIV2, the medium was replaced with fresh DMEM/F-12 plus 10% FBS, and clodronate-loaded liposomes (2.6 μg/mL; Liposoma BV, Netherlands) were added twice weekly to remove residual macrophages. Twelve to fourteen days later (DIV14-16), the culture medium was completely removed and replaced with either DMEM or Nb supplemented with 2% B27, 1% N2 and 1% antibiotics (referred to as DMEM^S^ and Nb^S^, respectively). After three additional days of culture, the DMEM^S^- and Nb^S^-based ACM were recovered, sterilized with 0.2µm syringe filters and frozen −20°C until use.

### Primary mouse midbrain culture

Primary mouse midbrain cultures were prepared as described previously [31]. Briefly, embryos (E13.5) from pregnant C57Bl6 females (Janvier LABS, Le Genest St Isle, France) were collected in L15 medium, and the midbrain was dissected under a binocular microscope. Midbrain tissue was digested in Trypsin plus 0.05% EDTA (Thermo Fisher Scientific, #25300054) for 20 min at +37°C and after neutralization of Trypsin with L15 supplemented with 10% fetal calf serum, predigested tissue was then mechanically triturated by gentle pipetting (8–10 strokes) using a Gilson pipette fitted with a sterile polypropylene blue tip with no filter (StarLab France, Orsay, France). After 10 min of sedimentation on ice, the supernatant was collected, and the trituration and sedimentation steps were repeated one more time. The collected supernatant was then centrifuged at 317xg for 5 min at +4°C. The cell pellet was resuspended in L15 medium and dissociated cells were distributed in PEI-coated Nunc 48-well plates (Roskilde, Denmark) in Nb at a density of 0.4 midbrains/well. After 1-2 hours incubation, ARA-C (1.2 µM; Sigma Aldrich, C1768) was added to curtail astrocyte proliferation. At DIV 1, ARA-C was brought to a 1.6 μM final concentration and at DIV 2, the culture medium was fully replaced by Nb^S^-based ACM. The use of a conditioning step by astrocytes was shown to improve the ability of Nb to promote the development and long-term viability of cultivated neurons [31, 32].

### Quantification of ROS levels

To determine the intracellular production of superoxide anions upon stimulation we used nitro-blue tetrazolium chloride (NBT) reagent following a protocol adapted from [33]. In brief, 100 000 cells were seeded in 48-well plates and stimulated as described above for 24 hours. One and a half hours before the assay, NBT (500 µM) was added to each well and the cells were further incubated before they were washed twice with warm PBS at 24 hours after stimulation. Then, 100 µL of absolute methanol was added to each well and the culture plates were left under the fume hood until methanol evaporation. After complete drying, 60 µL of 2 M KOH and 70 µL of DMSO were added to the wells and incubated under mild agitation for 5-10 minutes after the absorbance at 620 nm was measured. The results were normalized to those of the NSC experimental condition.

### Cytokine and glutamate assays

Cytokine levels were measured using ELISA kits or MSD U-Plex plates (Meso Scale Diagnostics) according to the manufacturer’s instructions. Glutamate levels were measured using an Amplex Red Glutamic Acid/Glutamate Oxidase Kit according to the manufacturer’s instructions. Quantification was carried out on 10 μl of culture medium samples. The fluorescent reaction products resulting from glutamic acid oxidation were quantified using excitation and emission wavelengths of 545 and 590 nm, respectively. Absorbance and fluorescence were measured using a SpectraMax i3X microplate reader (Molecular Devices, Sunnyvale, CA).

### Transcriptomic analysis

To determine the transcriptomic changes evoked by the different inflammatory stimuli, 1x10 microglial cells were seeded in 48-well plates and then stimulated or not (NSC) with LPS (10 ng/ml) or TPF for 24 hours. Seven to eight biological replicates per condition from two independent experiments were used. RNA was isolated using a NucleoSpin RNA XS Kit, and RNA quality was evaluated with an Agilent 2100 Bioanalyzer (Agilent Technologies). All RNA samples had RNA integrity numbers (RINs) greater than 7. We produced libraries for 31-mRNA sequencing using 200 ng of purified RNA per sample processed with a TruSeq Stranded RNA Kit (Illumina, San Diego, CA) according to the manufacturer’s protocol. Paired-end sequencing was subsequently performed on the Illumina NextSeq 500 and NovaSeq 6000 platforms with a sequencing depth of 80 million reads per sample (iGenSeq core facility, ICM, Paris France). All samples were assessed separately for overall read quality using FASTQC. Reads were trimmed using the Trimmomatic tool (Illumina), and then aligned to the mouse genome using Top-Hat (Illumina) and fragments were assigned to genes using the FeatureCounts program (iCONICS core facility, ICM, Paris France). Differentially expressed genes (DEGs) were analyzed using the open-source Bioconductor package DESeq2 in R software [34] (log2FC > 0.5; FDR-adjusted *p* value < 0.05). Pathway enrichment analysis was performed using the open-source gene enrichment analysis tool Enrichr [35-37]. Data visualization and biological interpretation were performed with GraphPad Prism 7.0 software.

### Metabolomic analysis

Metabolite profiling analysis was carried out on an Acquity UPLC system (Waters Corp, Saint-Quentin-en-Yvelines, France) coupled to a hybrid Orbitrap- and Q Exactive-based instrument (Thermo Fisher Scientific, Illkirch, France). To analyze and compare the microglial cell metabolome under different inflammatory conditions, we performed an untargeted metabolomic analysis of cells stimulated with or without LPS (10 ng/mL) or TPF^PD^ for 48 hours. Five biological replicates for each condition were prepared. Approximately 1x10^6^ cells grown in 25cm^3^ flasks were lysed using a methanol-based method consisting of adding 1 mL of frozen methanol (-20°C) followed by 5 min of incubation on dry ice and 5 min of rest at room temperature. This incubation cycle was repeated 3 times before the cells were collected with the help of a cell scraper, after which the cells were transferred to a 50 mL Falcon tube. The samples were kept frozen at -80°C until use. Sample preparation for LC-MS analysis was as follows: Prior to extraction, cells were homogenized in 100 µL of H_2_0 containing 0.1% formic acid and internal standards (a mixture of 16 labeled amino acids at 10 µg/mL) to a final concentration of 2500 cells/µl. Six cycles of freeze-thaw cycling were applied for cell lysis. Then, 4 volumes of frozen methanol (−20 °C) containing an equimolar (10 µg/mL) mixture of internal standard [16 labeled amino acids and 3 labeled nucleotides (ATP, N_5_; ADP, N_5_ and AMP, C, N_5_)] were added to 100 μL of sample and vortexed. The samples were further sonicated for 10 min before being centrifuged at 10,000×g and +4°C for 2 min and left for one hour (+4°C) for slow protein precipitation. The samples were then centrifuged at 20,000×g (+4 °C) for 20 min, after which the supernatants were transferred to new tubes and subsequently dried before being reconstituted in a solution of water/acetonitrile (40:60; v-v). The chromatographic conditions and experimental settings were previously described [38]. Briefly, LC-MS analysis was performed with a ZIC-*p*HILIC chromatographic column (5 µm, 2.1 × 150 mm, Merck, Darmstadt, Germany). Full scan positive and negative ionization modes with a resolution of 70.000 (FWHM) and a scan range of m/z 50–750 were used. The mass spectrometer was systematically calibrated in both ion polarity modes with Pierce calibration solution. The injection volume was set to 10 μL. The mobile phase (flow rate of 0.2 mL/min for 29 min) consisted of 10 mM ammonium carbonate at pH=10.5 (A) and acetonitrile (B). MS data processing was performed using the XCMS R package with the CentWave algorithm and CAMERA tools implemented in R software and the galaxy workflow4metabolomics [39-41]. The processing steps included peak picking, peak grouping, retention time correction and annotation of isotopes and adducts. Processed LC-MS data were further analyzed based on standard protocols [42, 43], which resulted in a data matrix in which each metabolomic feature was characterized by a retention time (RT), a mass-to-charge ratio (m/z), its corresponding intensities and the isotope-adduct annotation from the CAMERA tool. The metabolomics data matrix was then filtered, normalized, curated and log-10 transformed based on a quality assurance (QA) strategy [44, 45]. Notably, peaks with more than 30% of missing values were discarded. Data visualization for biological interpretation was performed with the Multi Experiment Viewer (MeV) statistical software package (version 4.9.0; http://www.tm4.org/mev/) [44]. This software was used to perform principal component analysis (PCA), heatmaps and tests to compute a p-value for each metabolic feature with a threshold of significance set to 0.05. False discovery rates (FDRs) were corrected using the Benjamini-Hochberg method [47] to adjust p-values for false discovery involving multiple comparisons.

### Quantitative real-time PCR

Total RNA was isolated using an extraction kit according to the manufacturer’s instructions (Macherey-Nagel). The RNA concentration was assayed with a NanoDrop (Thermo Scientific) and adjusted with nuclease-free water to an equal concentration for all samples. RNA was then reverse transcribed to cDNA using the Verso cDNA Synthesis Kit (Thermo Scientific) following the manufacturer’s instructions. In brief, RNA was denatured at 70°C for 2 min. Then, the kit mixture was added to each tube. The samples were subjected to successive incubations at 25°C for 10 min, 42°C for 1 h and 85°C for 5 min.

A quantitative PCR assay was designed to include three technical replicates per sample and two reference genes (*Rab7* and *Sdha* for primary mouse microglia; *Ubc* and *Sdha* for human microglia-like cells) which were selected upon analysis of gene expression stability across the different experimental conditions (geNorm tool of qBasePlus software, Biogazelle) and among the 8-10 initial gene candidates. Quantitative PCR was performed using a LightCycler® 480 System with compatible SYBR Green reagents (LightCycler® 480 SYBR Green I Master). The PCR program was 95°C for 10 min, 45 cycles of 95°C for 15 sec and 60°C for 60 sec and cooling. All the generated data were further processed and analyzed with qBasePlus.

### Generation of human induced microglia-like cells

Blood samples were collected from a healthy donor (49-year-old male) who provided signed informed consent and who was included in an INSERM-sponsored clinical study. Human induced microglia-like cells (iMGs) were obtained following the protocol published by [48] with some modifications. Briefly, 10 mL blood samples were collected in EDTA, and peripheral blood mononuclear cells (PBMCs) were purified by density gradient (Histopaque-1077) centrifugation at 400xg for 30 min. After isolation, the cells were washed in RPMI-1640 medium and centrifuged at 300xg for 10 min. The cell pellet was resuspended in separation buffer (MACS BSA stock solution/AutoMACS® Rinsing Solution at 1/20 dilution) and passed through MACS SmartStrainers (30 µm) to eliminate cell clumps. Monocytes were then isolated using 250 μL of MicroBeads-conjugated human CD14 antibodies (StraightFrom LRSC CD14 MicroBead Kit; Miltenyi Biotec) added to 40 mL of cell suspension. The cells and beads were incubated for 15 min at +4°C and centrifuged at 300xg for 10 min. The cell pellet was resuspended in the separation buffer and the labelled cells were separated by LS columns on a QuadroMACS™ Separator and recovered in whole-blood column elution buffer. After centrifugation at 300xg for 10 min, the cell pellet was resuspended in RPMI-1640 GlutaMAX supplemented with 10% FBS and 1% antibiotic-antimycotic solution. Monocytes were then seeded at a concentration of 6x10 cells/well in 96-well plates and incubated overnight at 37°C with 5% CO_2_. To induce differentiation, the medium was changed at DIV1 with RPMI-1640 GlutaMAX containing recombinant human GM-CSF (10 ng/ml) and recombinant human IL34 (100 ng/ml). The differentiating medium was freshly renewed at DIV6. On day 27 following cell seeding, differentiated iMGs were subjected to an inflammatory stimulation assay.

### Cell immunostaining and quantification

Primary microglia and iMGs were fixed with 3.7% formaldehyde solution (Sigma Aldrich, 252549) for 12 min and then washed twice with PBS. Primary mouse microglia were then incubated overnight at +4°C with rat anti-CD25 (1/100) or rat anti-CD11b (1/100) antibodies diluted in PBS. Then, cells were washed three times with PBS and incubated with anti-rat IgG coupled with Alexa488 fluorophore (1/300 dilution) for 2 hours at RT. For double immunofluorescence staining of iMGs, cells were incubated overnight at +4°C with rabbit anti-CX3CR1 and mouse anti-CCR2 primary antibodies (diluted 1/500 in PBS containing 0.1% Triton-X 100 and 5% NGS). The cells were then rinsed three times with PBS and incubated with anti-rabbit IgG conjugated to the Alexa488 fluorophore and anti-mouse IgG conjugated to the Alexa546 fluorophore (1/300 dilution) for 1 hour at RT. After two washes with PBS, the immunostained cells were further stained with Hoechst (1/4000 dilution) for 10 min and rinsed three additional times with PBS. The ratio of CX3CR1 to CCR2 was calculated by quantifying the fluorescent signal intensity using an automated quantitative imaging platform (ArrayScan, Thermo Fisher). Representative images were acquired using an Axio Observer 7/Apotome microscope (Zeiss). For midbrain cultures, a similar immunodetection protocol was used to assess tyrosine hydroxylase (TH), a dopaminergic neuronal marker. The cells were incubated at +4°C for 24 hours in mouse anti-TH primary antibodies (LNC1; Immunostar, #22941) diluted 1/2500 in PBS plus 0.2% Triton-X 100. The cells were then rinsed three times and incubated for 2 hours at room temperature in Alexa Fluor 488-conjugated goat anti-mouse secondary antibodies (1/300 dilution; Thermo Fischer Scientific). Following two washes in PBS, cells were stained with Hoechst 33342 (1/4000 dilution; Tocris, 5117) for 10 min. Neuron culture images were acquired with a Nikon Eclipse Ti-U fluorescence inverted microscope (Nikon France, Champigny sur Marne, France) equipped with a Hamamatsu’s ORCA-D2 camera and HCImage software (Hamamatsu Photonics, Massy, France). The number of TH+ neurons/culture well was estimated by visually inspecting samples with a 10× objective over 10–15 visual fields that were randomly selected for each treatment condition.

### Statistical analysis

Statistics were analyzed using SigmaPlot 14.0 or GraphPad Prism 7.0 software. The results are expressed as the means ± standard errors of the means (SEM). The normality of data distribution was assessed with the Shapiro-Wilk test. Statistical significance of normally distributed data was assessed using Student’s t-test for comparisons between two groups and ANOVA followed by Tukey’s post-hoc test for multiple comparisons. When the data were not normally distributed, significance was determined with Mann-Whitney U test for comparisons of two groups and Kruskal-Walli’s test followed by Dunn’s test for multiple comparisons. The statistical significance level was set as follows: * *p* value < 0.05, ** *p* value < 0.01, and *** p value < 0.001.

## Results

### Generation and characterization of patient-derived αSYN assemblies

To investigate the inflammatory properties of patient-derived αSYN aggregates we first generated pure *in vitro* amplified αSYN fibrillar assemblies using a protein misfolding cyclic amplification assay (PMCA) applied to LP-rich PD and DLB brain tissue homogenates as described elsewhere [15]. A detailed structural and biochemical characterization of this PMCA-amplified pathological material demonstrated that fibrillary assemblies derived from PD patients (F^PD^) are structurally different from those obtained from DLB patients (F^PD^). Thus, TEM analysis revealed disease-specific differences in the shape of the fibrils with PD patient-derived αSYN fibrils exhibiting a relatively flat and twisted appearance whereas those derived from DLB patients were cylindrical exhibiting no twists (Additional File 1: Fig. S1). Furthermore, assemblies from individual PD or DLB patients had distinct limited proteolysis patterns suggesting the existence of disease-specific αSYN strains [15].

### PD-derived **α**SYN assemblies evoke robust inflammatory response in primary microglial cells

We evaluated and compared the inflammatory potential of PD patient-derived αSYN fibrils by exposing primary microglial cells to equal concentrations (3µM) of either F^PD^ or well-characterized αSYN fibrils generated *de novo* (F^S^) [8, 10]. As a positive control for inflammatory microglia, cells were treated with LPS (10 ng/ml), a prototypical inflammogen and ligand for TLR4 [49]. After 48 hours of treatment, stimulated microglia undergo morphological changes as compared to non-stimulated control cells (NSC) (Fig. 1A). In particular, LPS-exposed cells lose their ramified and spindle-shaped processes adopting more flattened and wider morphologies typical of inflammatory induction [30]. Of note, F^S^ and F^PD^ treatment induced morphological changes similar to those observed with LPS stimulation suggesting inflammatory activation by αSYN fibrillar assemblies. To obtain further insights into the inflammatory potential of F^PD^, we evaluated the production and release of several inflammatory cytokines in response to increasing amounts of F^PD^ ranging from 0.01 to 3 µM. We observed a dose-dependent increase in the release of TNFα, IL6 and IL10 after 24 hours of treatment (Fig. 1B), with efficient and significant cytokine release occurring at F^PD^ concentrations of 1 µM and above (Fig. 1B). As inflammatory microglia, particularly when exposed to fibrillary aggregates of αSYN, can release large amounts of glutamate [50], we measured glutamate concentrations in microglial cell culture medium after exposure to F^PD^. As shown in Figure 1B, F^PD^ strongly promoted glutamate release from microglia in a dose-dependent manner within the F^PD^ concentration range of 1.5 and 3 µM (3- and 11-fold increases, respectively, compared to those of NSCs). Whereas both F^PD^ and F^S^ efficiently triggered inflammatory-associated morphological changes in microglial cells (Fig. 1A), we found that F^PD^ was more potent than F^S^ at inducing cytokine and glutamate release after 48 hours of stimulation. Indeed, at an equal concentration of αSYN assemblies (3µM), F^PD^ induced 5-, 17- and 3-fold more TNFα, IL6 and IL10 than F^S^, respectively (Fig. 1C). Similarly, the extracellular amount of glutamate following exposure to F^PD^ was twice as high as that in F^S^-stimulated cells (Fig. 1C). Interestingly, we observed that microglia-associated inflammatory responsiveness toward F^DLB^ was even more important than that evoked by F^PD^ assemblies (Fig. 1C). Together, our data not only show that F^PD^ has potent inflammatory effects on microglial cells but also that PD-derived assemblies exhibit stronger and weaker stimulatory effects than αSYN fibrils generated *de novo* and from F^DLB^, respectively. Given the structural differences between F^PD^ and F^S^, and between F^PD^ and F^DLB^ [10, 15], our data also suggest a structure-function relationship regarding the inflammatory properties of αSYN aggregates on brain tissue macrophages and argue for the use of PD-derived material to better model disease-associated inflammatory mechanisms.

**Figure 1.**
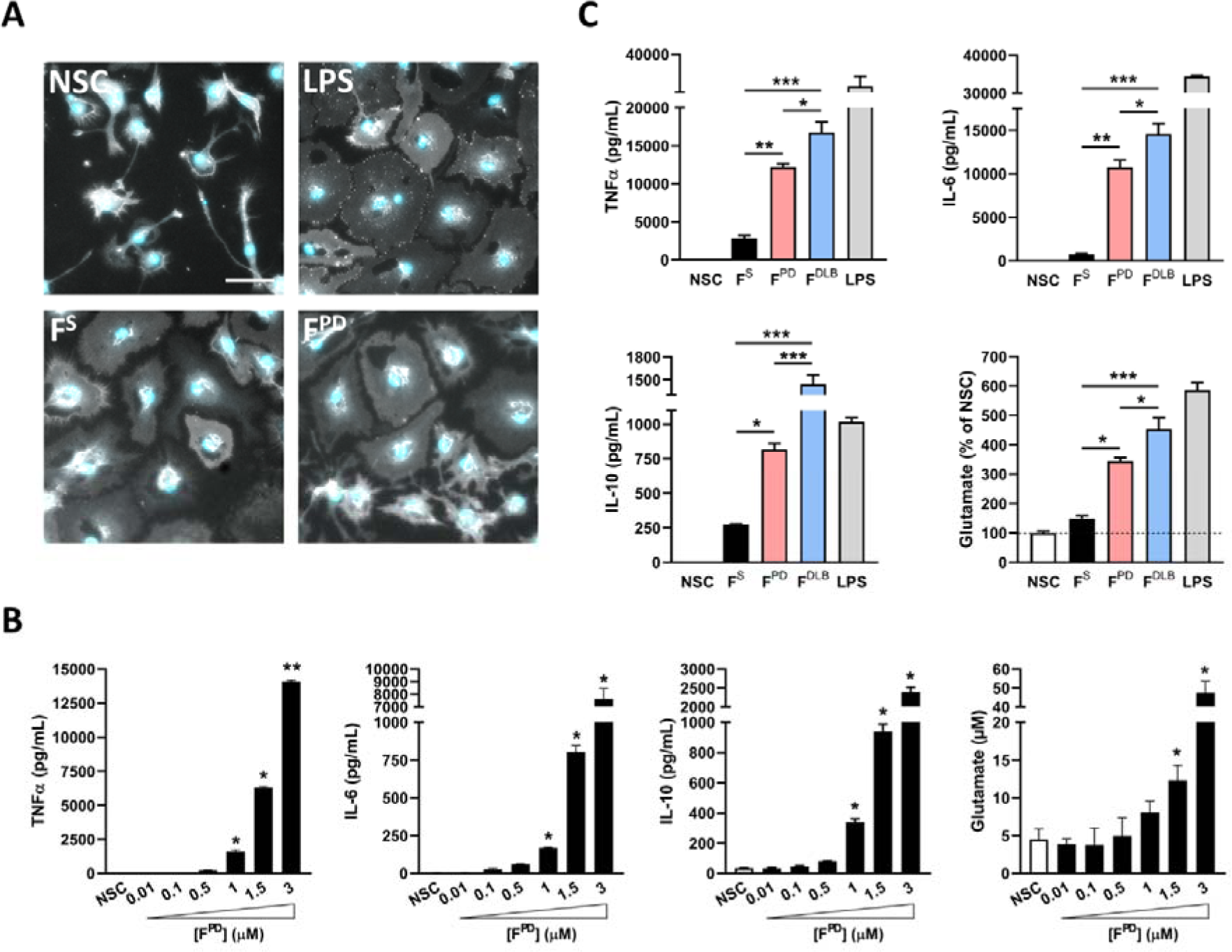
aSYN fibrils derived from PD patients hold more potent inflammatory properties than *de novo*-generated counterparts. (**A**) Representative images of CD11b immunostaining and Hoechst nuclear staining (blue) showing the morphological changes in microglial cells following inflammatory stimulation by the prototypal inflammogen LPS (10 ng/ml), *de novo* assembled human αSYN fibrils (3 µM; F^S^) and fibrils derived from PD patients (3 µM; F^PD^). Note that inflammatory stimulation results in cellular flattening and shortening of cell processes under all three inflammatory conditions. Scale bar: 50 µm. (**B**) Dose-response analysis of TNFα, IL6, IL10 and glutamate release by microglial cells exposed or not (NSC) to increasing concentrations (from 0.01 to 3 µM) of αSYN fibrils derived from PD patients (F^PD^). The data are presented as the means ± SEM (n = 6). \**p* < 0.05; \*\**p* < 0.01 vs. NSC (Tukey’s test). (**C**) Quantification of TNFα, IL6, IL10 and glutamate release by microglial cells exposed or not (NSC) to similar concentrations of F^S^, F^PD^ or F^DLB^ (3 µM). The data are presented as the means ± SEM (n = 6). \**p* < 0.05; \*\**p* < 0.01; \*\*\**p* < 0.001 vs. F^S^ (Tukey’s test).

### Chronic-type stimulation of microglia is associated with decreased cytokine levels but increased glutamate release

Chronic inflammatory conditions in humans are associated with sustained high tissue levels of TNFα, PGE_2_ and innate immune receptor-activating ligands that act in concert to shape the macrophage response. Combinatory stimulation of human macrophages with these factors (the so-called TPP stimulation) results in a specific molecular pattern and phenotype that departs from the well-established classical M1 and alternative M2 activation state axis [21]. In particular, the cell-surface markers CD25, CD14 and CD23 as well as the cytokine IL1α and, to a lesser extent, the chemokine CXCL5 were found to be induced in TPP-specific conditions [21]. Remarkably, elevated levels of both TNFα and PGE_2_ have been documented in the *substantia nigra* of PD patients, suggesting that these inflammatory mediators might act in concert with TLR-activating pathological αSYN aggregates to drive microglial cell polarization toward a specific disease phenotype. Therefore, in an effort to better model chronic-type PD innate immune response, we sought to integrate these disease-relevant inflammatory cues together with αSYN fibrils, the presumed primary pathological trigger. We first tested whether microglial cells would respond similarly to peripheral macrophages upon chronic-type inflammatory stimulation as previously described [21]. Indeed, while peripheral macrophages and microglia share an origin in the yolk sac, microglia are an ontogenetically distinct cell population, so their responsiveness to chronic-type inflammatory stimulation may not exactly mirror that of their macrophage relatives [51]. We thus investigated the expression of TPP-induced immune markers in microglial cells exposed to a combination of TNFα (800 UI/ml), PGE_2_ (1 µg/ml) and the TLR2 agonist Pam3CSK4 (Pam3C; 1 µg/ml) compared to cells stimulated with conditions linked to M1 (IFNγ; 20 ng/mL) or M2 (IL4; 10 ng/mL) polarization. Immunostaining for CD25 revealed that, unlike peripheral macrophages, microglial cells not only express high basal levels of this cell-surface marker but also exhibit unchanged expression under M2 and TPP stimulation, and downregulation under M1 polarization (Fig. 2A). Similarly, whereas IL1α was specifically induced in TPP-stimulated macrophages [21], it was only increased in M2-polarized (IL4-treated) microglial cells (Fig. 2B). Finally, TPP-exposed microglia produced and released as much CXCL5 as did IL4-treated cells, in contrast to the upregulation of this chemokine in both IFNγ- and TPP-stimulated human macrophages (Fig. 2B). Notably, the poor or degraded biological activity of recombinant IFNγ was unlikely to account for the apparent unresponsiveness of microglial cells to IFNγ-induced CXCL5 since this treatment efficiently stimulated the intracellular production of reactive oxygen species (ROS) as a consequence of NADPH-oxidase activation (Fig. 2C). Taken together, these data show that microglial cells feature distinct TPP-associated gene regulation patterns compared to those of peripheral macrophages.

**Figure 2.**
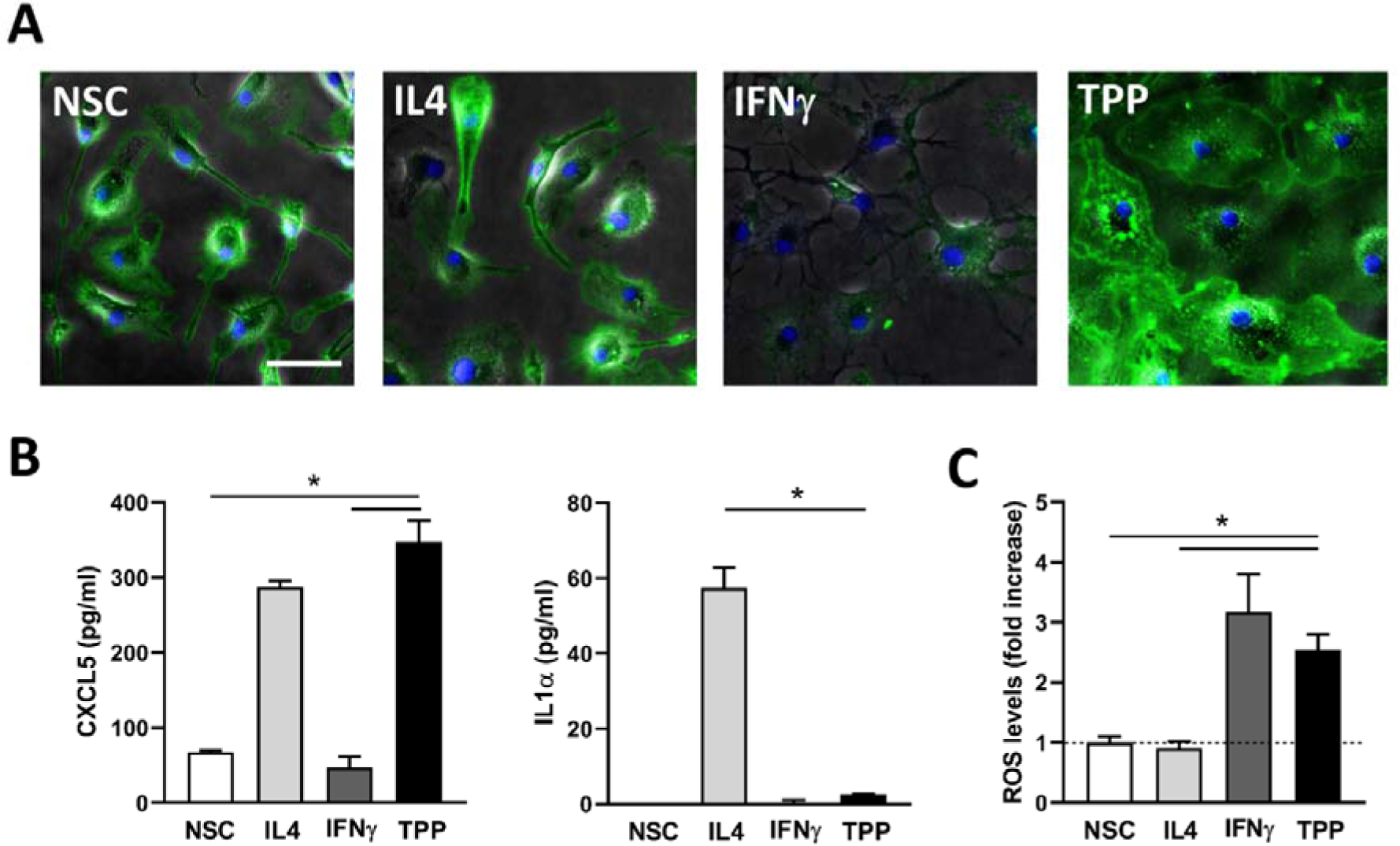
Regulation of human macrophage-associated TPP-specific inflammatory markers in microglial cells. (**A**) Representative images showing CD25 immunostaining with Hoechst nuclear stain (blue) in nonstimulated microglial cells (NSC) and cells exposed to IL4 (10 ng/mL), IFNγ (20 ng/mL) or a combination of TNFα (800 UI/ml), PGE_2_ (1 µg/ml) and Pam3C (1 µg/ml) (TPP stimulation). Scale bar: 60µm. (**B**) Quantification of CXCL5 and IL1α release by microglial cells exposed or not (NSC) to the indicated treatments. Data are means ± SEM (*n* = 4). \**p* < 0.05 vs. TPP (Tukey’s test). (**C**) Quantification of ROS production measured via the NBT reaction in microglial cells exposed or not (NSC) to the indicated treatments. The data are represented as the means ± SEM (*n* = 4). \**p* < 0.05 vs. TPP (Tukey’s test).

We next investigated the microglial cell response to chronic-type inflammatory stimulation by implementing PD patients-derived αSYN fibrils (F^PD^) to the treatment cocktail assuming that F^PD^ would efficiently bind and activate microglial TLRs as demonstrated for αSYN fibrils made *de novo* [8, 52-55]. Using ELISA assay, we found that unlike LPS and F^PD^, TNFα and PGE_2_, alone or in combination (TP), were ineffective at stimulating the release of TNFα, IL6, IL1β and IL10 after 48 hours of treatment (Fig. 3A and B). Remarkably, cotreatment of F^PD^-exposed cells with TNFα and PGE_2_ (TPF^PD^ stimulation) significantly suppressed cytokine released compared to that in cells treated with F^PD^ alone, suggesting that chronic-type inflammatory stimulation is associated with a significantly but less intensive inflammatory state. To determine whether TPF^PD^ stimulation is associated with an overall reduction in the inflammatory grade of microglial cells, we measured and compared intracellular ROS levels following the different treatments. Figure 3C shows that, unlike its effects on cytokine production/release, TP did not mitigate F^PD^-induced ROS generation in microglial cells. To further investigate the inflammatory properties associated with TPF stimulation, we analyzed microglial glutamate release under these conditions. We observed that, unlike cytokine (decrease) and ROS (unchanged) production, TPF^PD^ treatment was associated with a significant increase in extracellular glutamate levels compared to F^PD^ treatment alone (Fig. 3D). Overall, our data show that, upon chronic-type inflammatory stimulation, microglial cells adopt a nonconventional phenotype rather than the classical proinflammatory M1 state.

**Figure 3.**
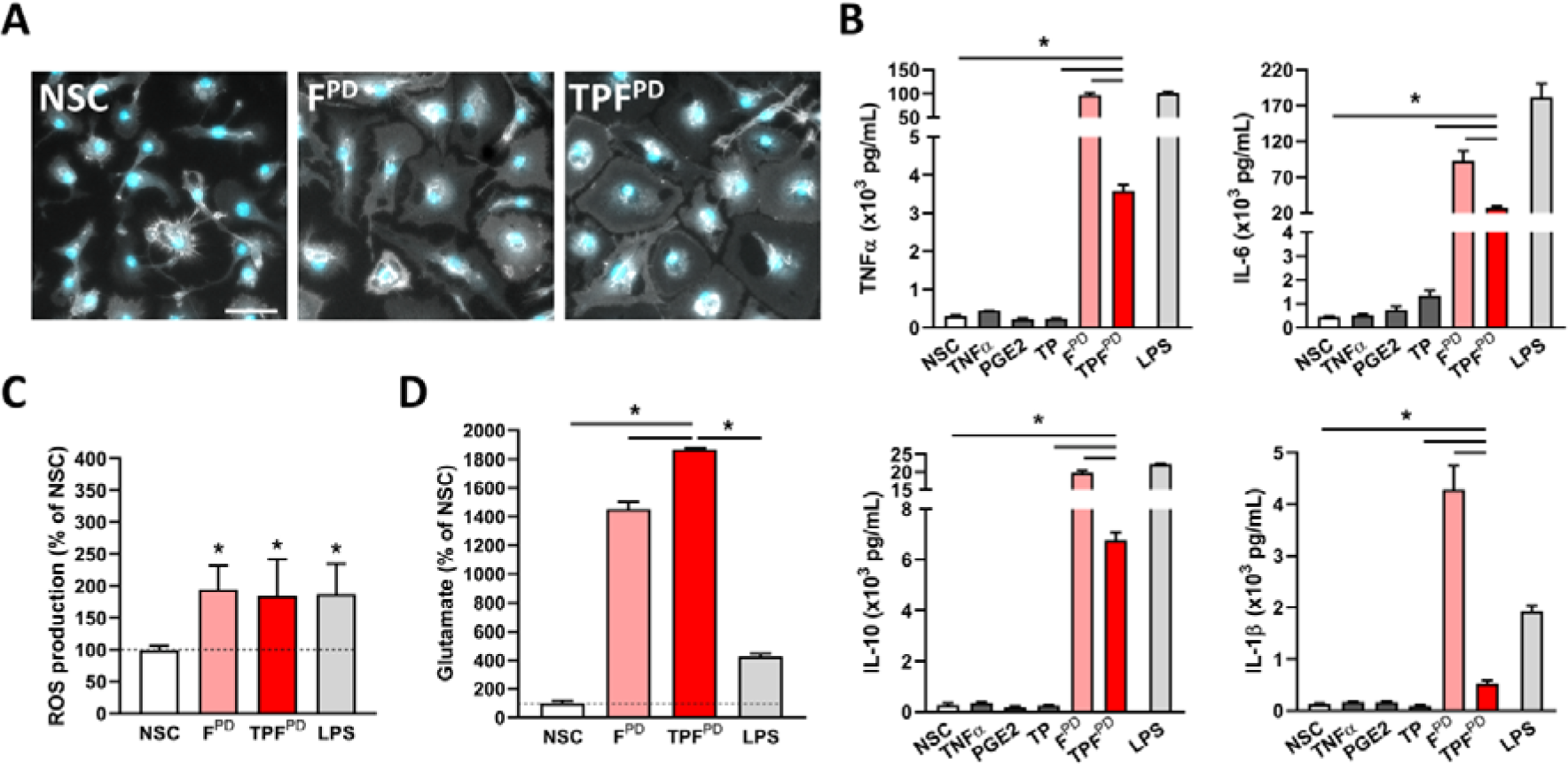
TPF^PD^-related chronic-type inflammatory stimulation of microglia is associated with lower cytokine but increased glutamate release. (**A**) Representative images of CD11b immunostaining and Hoechst nuclear staining (blue) of microglial cells following inflammatory stimulation by PD patient-derived αSYN fibrils (1.5 µM; F^PD^) or TPF^PD^ as compared to nonstimulated cells (NSC). Note that F^PD^ and TPF^PD^ induce similar morphological changes of microglial cells. Scale bar: 50µm. (**B**) Quantification of TNFα, IL6, IL10 and IL1β release by microglial cells exposed or not (NSC) to TNFα (800 IU/mL), PGE_2_ (1 µg/mL), TNFα+PGE_2_ (TP), F^PD^ (1.5 µM), TNFα+PGE_2_+F^PD^ (TPF^PD^) or LPS (10 ng/mL). The data are represented as the means ± SEM (n = 3-6). \**p* < 0.05; \*\**p* < 0.01; \*\*\**p* < 0.001 vs. F^S^ (Student’s t-test). (**C**) Quantification of ROS production measured by the NBT reaction in microglial cells exposed or not (NSC) to F^PD^ (1.5 µM), TPF or LPS (10 ng/mL). The data are expressed as a % of the NSC control. The bars are the means ± SEM (*n* = 6). \**p* < 0.05 vs. NSC (Student’s t-test). (**D**) Quantification of glutamate release by microglial cells exposed or not (NSC) to F^PD^ (1.5 µM), TPF^PD^ or LPS (10ng/mL). The data are expressed as a % of the NSC control. The bars are the means ± SEM (n = 3-6). \**p* < 0.05 vs. TPF^PD^ (Tukey’s test).

### Chronic-type inflammatory stimulation of microglia is associated with a specific genomic signature

The above data showing specific inflammatory features of microglia subjected to TPF^PD^ stimulation might indicate unique molecular pathway engagement and transcriptional programming as previously exemplified in peripheral macrophages [21]. To elucidate the molecular signature of this inflammatory polarization, we conducted unbiased transcriptomic analysis through RNA-sequencing (RNA-seq) performed on nonstimulated cells (NSCs), LPS-polarized cells as a reference for the M1 proinflammatory state, and TPF^PD^-activated microglia. To examine the effect of stimulation, we initially performed unsupervised principal component analysis (PCA). As expected, the samples were separated well by activation status (NSC vs stimulated) on the first principal component but also by type of stimulation (LPS vs TPF^PD^), which drove the second component of the variability (Fig. 4A). By using differentially expressed genes (DEGs; log2FC > 0.5, false discovery rate [FDR] adjusted *P* value < 0.05) between TPF^PD^, M1 (LPS) and NSCs we identified 554 upregulated and 456 downregulated genes in TPF^PD^-exposed cells. Unsupervised cluster analysis of the top 413 DEGs (by *P*-value) revealed a striking difference between the transcriptomes of M1-polarized and TPF^PD^-treated microglia (Fig. 4B). To obtain insights into how differential gene regulation translates into pathway modulation we first processed expression data with the Pathifier algorithm, which transforms gene-level information into pathway-level records [56]. Principal component analysis of the newly generated pathway- related dataset indicated that both the stimulation and type of stimuli drove strong pathway variability (Fig. 4C). To document in detail the pathways and biological functions altered by chronic-type inflammatory stimulation, we performed pathway enrichment (KEGG) and gene ontology (GO) term analyses. We found that among the most significantly enriched KEGG pathways in TPF^PD^- stimulated cells were ribosome biogenesis, glutathione metabolism, steroid biosynthesis, spliceosome, ferroptosis and RNA transport (Fig. 4D). The results from GO analysis confirmed the pathway enrichment data for the most part. Yet, they also revealed additional features related to the regulation of the inflammatory response and primary metabolic and lipid processes (Additional File 1: Fig. S2). Notably, increased glutathione metabolism in TPF^PD^-stimulated microglial cells might be possible under conditions of high cysteine supply and glutamate levels provided that the expression and/or activity of the rate-limiting enzymes glutathione cysteine ligase (GCL) and glutathione synthase (GSS) increase. One possible mechanism that could increase microglial cysteine levels upon inflammatory stimulation is the activation of xCT, a transmembrane cystine-glutamate exchange transporter encoded by the *Slc7a11* gene [57]. Remarkably, the *Slc7a11*, *Gss* and *Gcl* genes (*Gclc* and *Gclm*) were among the most upregulated genes in TPF^PD^-stimulated cells (Fig. 4E). Increased *Slc7a11* gene expression in TPF^PD^-stimulated microglia was consistent with the increased capacity of these cells to release glutamate (Fig. 3C).

**Figure 4.**
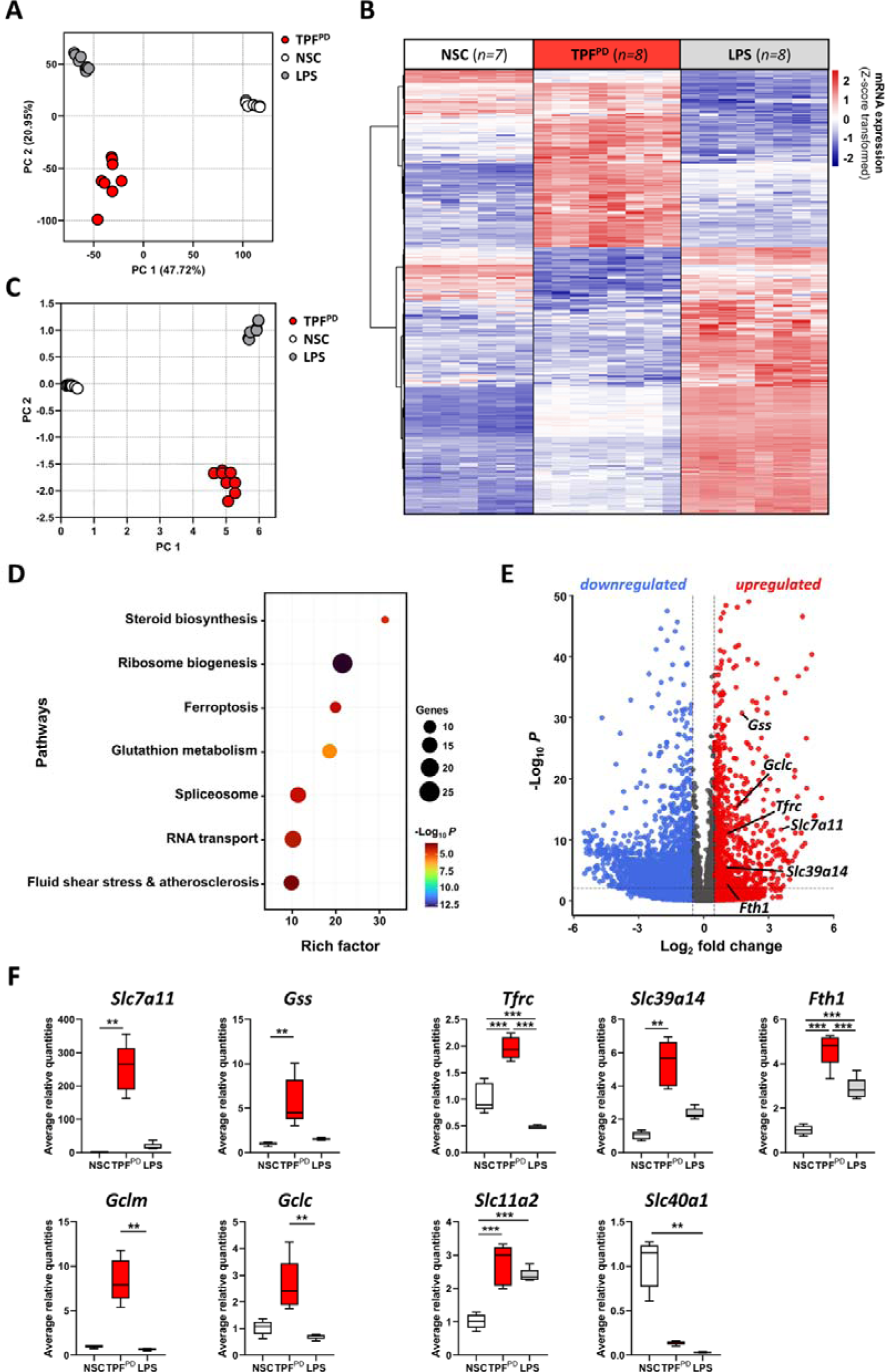
Specific transcriptional reprogramming of microglial cells upon TPF^PD^ chronic-type inflammatory activation. (**A**) Principal component analysis (PCA) of gene expression data based on the first two PCs shows, on the one hand, a clear distinction between nonstimulated (NSC) and activated cells (LPS and TPF^PD^) and, on the other hand, strong differences between M1-type (LPS) and chronic-type (TPF^PD^) inflammatory activation. (**B**) Hierarchical clustering and heatmap of differentially expressed genes (DEGs; n=413) in TPF^PD^-activated microglia versus nonstimulated (NSC) and LPS-treated cells. The scaled expression value (Z-score transformed) is shown in a blue-red color scheme with red indicating higher expression, and blue lower expression. Biological replicates are indicated in brackets. (**C**) PCA analysis of pathway-related data generated by the Pathifier method (Drier et al., 2013) demonstrates clear clustering and separation of stimulated and nonstimulated (NSC) cells and of M1-type (LPS) and chronic-type (TPF^PD^) inflammatory activation. (**D**) Bubble chart showing the enrichment of the KEGG Pathway in TPF^PD^-treated microglial cells (adjusted *p* < 0.05). The bubble size indicates the number of genes annotated in the indicated KEGG pathway. The colors represent pathway enrichment (% of overlapping genes) in TPF^PD^-exposed cells. (**E**) Volcano plot depicting individual DEGs (|Log_2_ Fold Change| > 2) in TPF^PD^-treated microglia versus M1-type (LPS) activated cells. The red and blue dots show upregulated and downregulated genes in TPF^PD^-treated cells, respectively. Individual genes of interest are indicated. (**F**) Relative gene expression analysis (qPCR) of individual gene candidates in nonstimulated-(NSC), LPS-activated and TPF^PD^-exposed microglial cells. The data are represented as the means ± SEM (n=5 biological replicates from independent experiment). \**p* < 0.05 vs TPF^PD^ (Student’s t test).

Another interesting observation is the apparent link between TPF^PD^ stimulation and ferroptosis. Ferroptosis is a type of redox-driven programmed cell death process distinct from classical apoptosis. This process might occur because of uncontrolled iron metabolism, lipid peroxidation and thiol regulation, three processes that are commonly found in the proinflammatory environment. Since TPF^PD^-stimulation does not induce microglial cell death under our experimental conditions (Additional File 1: Fig. S3), it is likely that genes involved in iron metabolism contributed to the ferroptosis term in our pathway enrichment list. Indeed, we found that iron transport- and storage-related genes, including *Tfrc*, *Slc39a14* and *Fth1* (encoding for the Transferrin receptor, the zinc and non-transferrin-bound iron transporter Zip14, and the Ferritin heavy chain, respectively) were strongly and differentially upregulated in TPF^PD^-treated cells (Fig. 4E). The genes encoding divalent metal transporter 1 (DMT1) and the iron extruder Ferroportin (*Slc11a2* and *Slc40a1*) were similarly expressed in LPS- and TPF^PD^-exposed cells (Fig. 4E). Overall, our data suggest differences in microglial iron handling under two distinct inflammatory conditions.

To validate our transcriptomic data, we performed qPCR expression analysis of gene candidates using total RNA isolated from microglial cells exposed or not to LPS and TPF^PD^ in an independent experiment. We confirmed the differential expression of *Gss*, *Gclc, Gclm* and *Slc7a11* in TPF^PD^-treated cells compared to that in nonstimulated and LPS-exposed microglia (Fig. 4F). Likewise, iron metabolism-related genes (*Tfrc*, *Slc39a14*, *Fth1*, *Slc11a2* and *Slc40a1*) exhibited similar changes in expression as identified via transcriptomic analysis (Fig. 4F). Overall, we identified genes and pathways that define microglia polarized upon TPF^PD^ chronic-type inflammatory conditions.

### TPF**^PD^**-related chronic-type inflammatory stimulation of microglia is associated with M1-related metabolic changes

The above transcriptome-based analysis revealed pathway enrichment linked to metabolic processes. It is now well established that immune activation rapidly and substantially enhances metabolic outputs that in turn regulate immune responses [58]. For instance, in macrophages, including microglia, M1 phenotype polarization is accompanied by a shift from oxidative phosphorylation to aerobic glycolysis for energy production [59]. To further investigate TPF^PD^- associated metabolic changes in microglia, we performed untargeted metabolomic analysis under the same experimental conditions as above.

Our results revealed significant alterations in the levels of 248 metabolites in TPF^PD^-treated cells. Indeed, hierarchical clustering analysis of metabolite datasets and heatmap generation (Fig. 5A) showed important differences between nonstimulated cells and microglia exposed to either LPS or TPF^PD^. The metabolome profile of TPF^PD^-activated cells displayed only subtle differences from that of LPS-treated microglia. Of the 248 metabolites showing significant changes, only 28 were annotated in the databases. Unsupervised cluster analysis of these 28 known metabolites confirmed the close relationship between the two inflammatory conditions (Fig. 5B). In particular, both stimulations resulted in decreased α-ketoglutarate/succinate and fumarate/succinate ratios, which are indicative of tricarboxylic acid (TCA) cycle breaks and metabolic switches from oxidative phosphorylation to glycolysis, as previously reported [60]. Hence, these data suggest that from a respiratory point of view, TPF^PD^-polarized microglia behave as classical proinflammatory M1 cells. Moreover, although TPF^PD^-polarized microglia release more glutamate than LPS-activated cells (Fig. 3C), these two inflammatory stimulations are characterized by comparable increases in glutamate and glutathione disulfide levels (Fig. 5C). Hence, our data showed that metabolic changes associated with M1 and TPF^PD^ microglial polarization are closely related.

**Figure 5.**
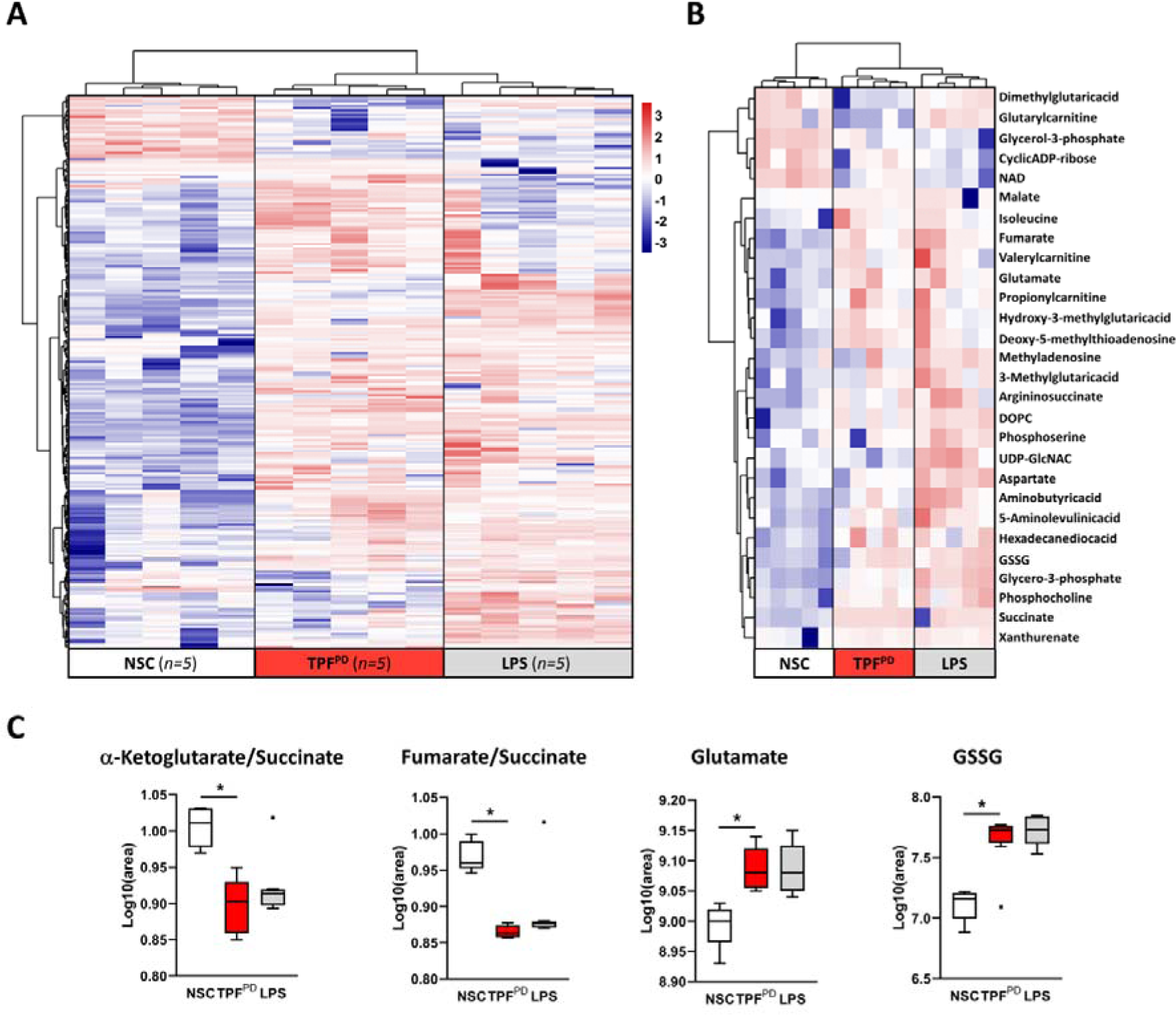
Microglial metabolic reprogramming evoked by TPF^PD^ is closely related to M1-type polarized cells. (**A**) Hierarchical clustering analysis and heatmap visualization of 248 differentially regulated metabolites (28 annotated and 220 unknown/nonannotated metabolites) in stimulated cells (LPS and TPF^PD^) compared to those in the NSC control group. The number of biological replicates is indicated in brackets. Clear changes in the metabolic profile between stimulated and nonstimulated cells are observed. (**B**) Hierarchical clustering analysis and heatmap visualization of the 28 annotated metabolites extracted from (A). Despite these subtle differences, the TPF^PD^-and LPS-associated metabolic signatures are closely related. (**C**) Box plots showing the relative amount of a-ketoglutarate, succinate, fumarate, glutamate and glutathione disulfide (GSSG) in control (NSC) and in LPS- and TPF^PD^-treated microglial cells. Similar decreases in the a-ketoglutarate/succinate and fumarate/succinate ratios were found in LPS- and TPF^PD^-stimulated cells. Likewise, similar increases in glutamate and GSSG levels were observed under both stimulatory conditions. \**p* < 0.05 (Kruskal-Wallis & Wilcoxon test).

### Conserved TPF **^PD^**-specific induction of Tfrc and Slc7a11 gene expression in human induced microglial-like cells

Integration and comparison of human and murine transcriptome datasets indicate both divergent and conserved gene expression regulation upon inflammatory stimulation between species [61]. Our comparative study of TPP-induced inflammatory markers in mouse microglial cells clearly illustrates such divergence indicating cell-type and/or species specificity (Fig. 2). To further determine the relevance of our data generated from mouse microglia, we prepared and used induced-microglia-like cells (iMGs) differentiated from human monocytes isolated from healthy donors [48]. Successful differentiation of monocytes into iMGs was confirmed by investigating the expression of the CCR2 and CX3CR1 chemokine receptors. As shown in Figure 6A, monocyte differentiation into iMGs was accompanied by a significant increase in the CX3CR1/CCR2 ratio indicating that differentiated cells acquired a macrophage-/microglia-like phenotype [60]. After 24 days of differentiation, iMGs were exposed or not (NSC) to LPS, F or TPF at concentrations equivalent to those used for mouse microglia experiments and then collected after 48 hours of activation for assessment by qPCR of candidate genes. We found that, similar to mouse microglial cells, TPF chronic-type inflammatory stimulation of iMGs resulted in specific upregulation of *Tfrc* and *Slc7a11* compared to that in response to LPS or F treatment (Fig. 6B). Likewise, although not significantly different, a clear trend toward increased expression of *Slc11a2* was observed under both LPS and TPF^PD^ conditions. In contrast, *Gclm* was found to be equally induced in LPS- and TPF^PD^-exposed cells but not in cells stimulated with F^PD^ alone. Interestingly, compared with LPS and F^PD^, TPF^PD^ stimulation specifically induced marked upregulation of *Tlr2* in iMGs in contrast to what we observed in mouse microglial cells. Moreover, while *Tlr4* expression was reduced upon LPS and F^PD^ treatment compared to that in control cells (NSC), it remained unchanged under TPF^PD^ conditions suggesting that the regulation of TLR expression in human microglia highly depends on the type of inflammation and inflammatory stimuli. Together, our data support previous findings showing species-dependent gene regulation in macrophages and demonstrate conserved TPF^PD^-associated *Tfrc* and *Slc7a11* gene regulation between mouse and human microglia. They further highlight that TPF^PD^-stimulated microglia may acquire a unique phenotype characterized by increased iron retention capacity and glutamate release.

**Figure 6.**
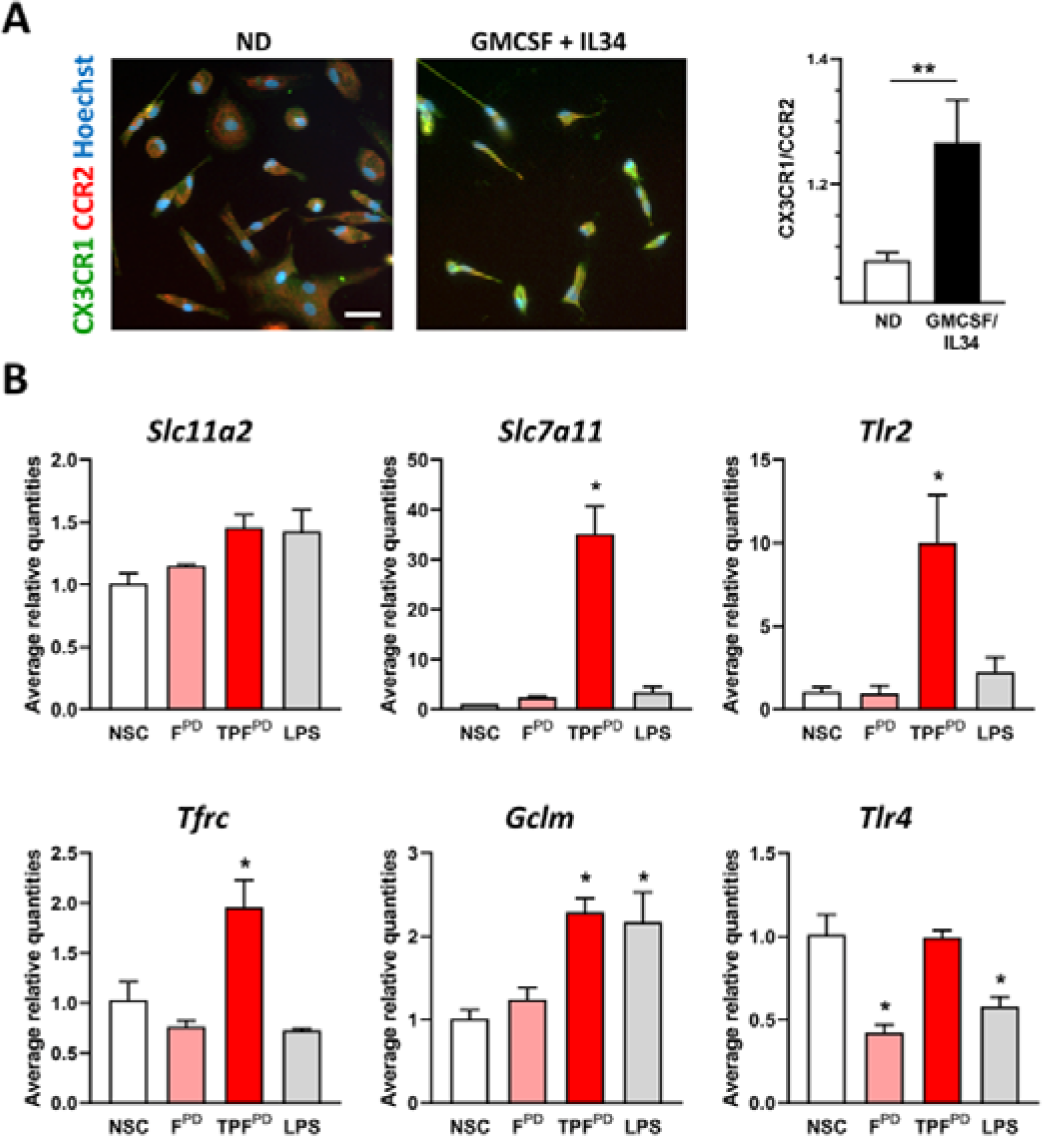
Compared with mouse microglia, TPF ^PD^-exposed human microglial-like cells exhibit both conserved and distinct gene regulation. (**A**) Representative images showing CCR2 (red) and CX3CR1 (green) double immunostaining with Hoechst nuclear stain (blue) in nondifferentiated (ND) and GM-CSF/IL34-differentiated human monocytes (iMG) after 16 days of culture. The right panel shows the quantification of the immunosignal CX3CR1/CRR2 ratio in ND and GM-CSF/IL34-differentiated cells. The data are presented as the means ± SEM (*n* = 4-5). \*\**p* < 0.01 vs. ND (Student’s t-test). Scale bar: 40 µm (**B**) Relative expression analysis (qPCR) of individual gene candidates in nonstimulated (NSC), LPS-, F^PD^- and TPF^PD^-activated iMGs. The data are presented as the means ± SEM (biological replicates n=3). \**p* < 0.05 vs. TPF^PD^ (Student’s t test).

### Chronic-type inflammatory stimulation of microglia results in xCT-dependent neurotoxicity on dopaminergic neurons

Our molecular and phenotypic analysis of microglial cells stimulated by chronic-type inflammatory cues revealed that cells under such activation are more prone to release high amounts of glutamate than classically activated (M1) proinflammatory microglia. This peculiar effector function of TPF^PD^- activated cells may be detrimental to dopaminergic neurons, which are particularly vulnerable to glutamate-induced excitotoxicity [63]. We sought to test this hypothesis using an experimental paradigm, allowing us to evaluate the neurotoxic potential of microglia-conditioned medium (MCM) on dopaminergic neurons. Specifically, we tested and compared the survival of dopaminergic neurons maintained in Nb^S^-based ACM, 24 hours after they received MCM (30%, v/v) from F^PD^- and TPF^PD^-stimulated microglia cultivated in DMEM^S^-based ACM (Fig. 7A). To prevent any possible direct neuronal effects of inflammatory stimulating factors (F^PD^ or TPF^PD^), we implemented a washout step 6 h after initiating microglial stimulation and maintained activated microglial cells for an additional 24 hours in fresh DMEM^S^-based ACM (Fig. 7A). At this time, MCM from F^PD^- or TPF^PD^-stimulated microglia displayed higher glutamate levels than did those from NSC (Fig. 7B). Under such experimental setup, we found that TPF^PD^-stimulated MCM induced significantly more TH+ dopaminergic neuron loss than did F^PD^-stimulated cells suggesting that chronic-type inflammatory stimulation drives microglia toward a more aggressive phenotype than classically activated M1 cells (Fig. 7C and D). Most interestingly, we observed that the rate of TH+ neuronal loss was strongly correlated with the glutamate content in the MCM suggesting that xCT-dependent glutamate release and glutamate-induced excitotoxicity may underlie key neurotoxic mechanisms supported by chronic-type inflammatory microglial cells (Fig. 7E). Importantly, the transfer of DMEM^S^-based ACM from control microglial cultures to midbrain cells had no impact on dopaminergic neuron survival.

**Figure 7.**
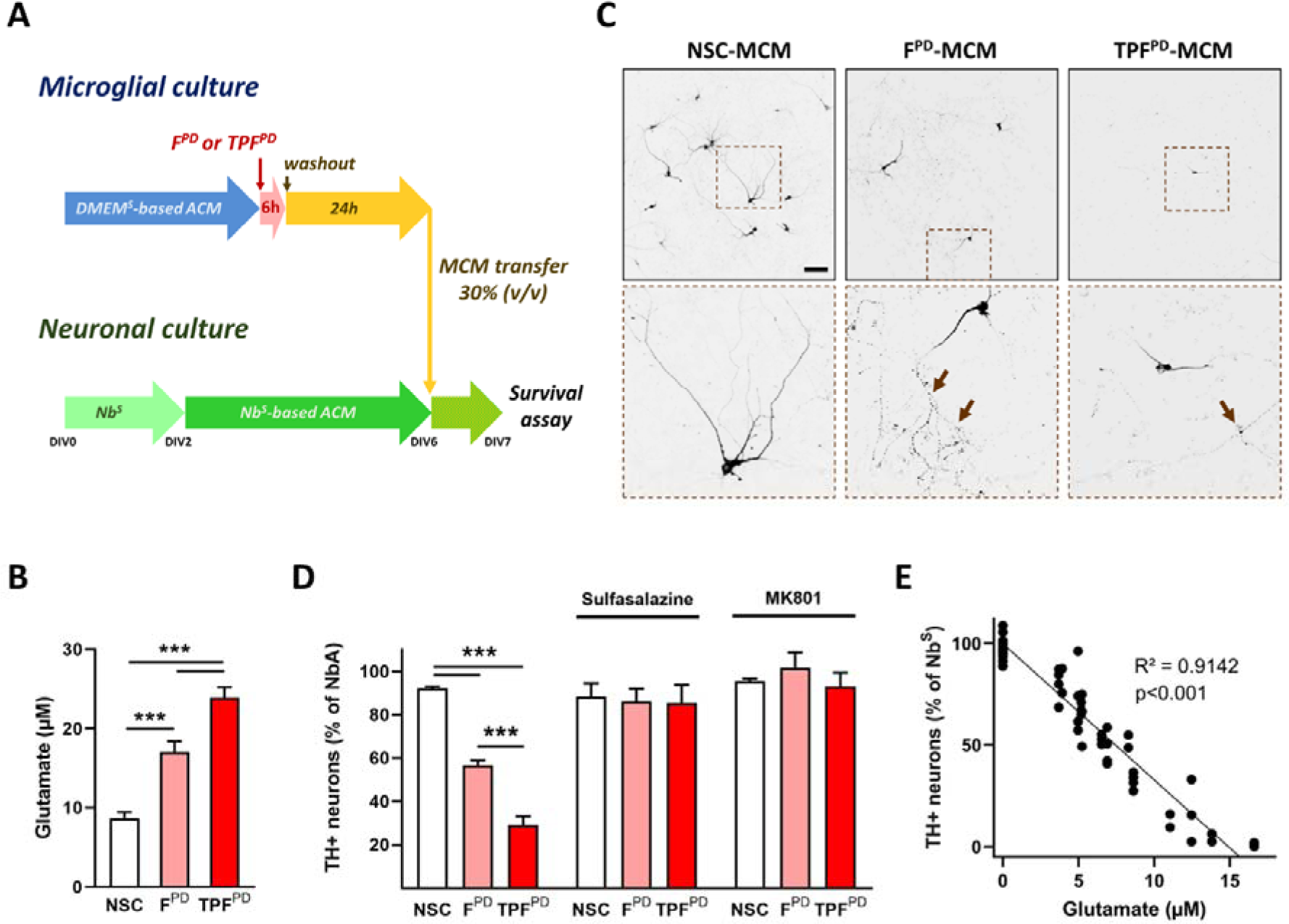
TPF^PD^-stimulated microglial cells trigger more excitotoxic cell death than F^PD^-exposed cells on dopaminergic neurons. (**A**) Schematic representation of the experimental setup used to analyze the neurotoxic potential of chronic-type inflammatory microglia. Microglial cell cultures were maintained in DMEM^S^-based astrocyte-conditioned medium (ACM) and exposed or not (NSC) for 6 hours to either F^PD^ or TPF^PD^. Following inflammatory stimulation, the culture medium was fully replaced with fresh DMEM^S^-based ACM (washout step), and the microglial cells were left for 24 hours before the glutamate assay (**B**). Midbrain neuronal cultures were prepared from E13.5 mouse embryos and maintained for 2 days in Nb^S^. On day *in vitro* 2 (DIV2), the culture medium was fully replaced with Nb^S^-based ACM and the neurons were left to mature until DIV6. At DIV6, 30% (v/v) of the Nb^S^-based ACM media was replaced with microglia-conditioned media (MCM) and TH+ dopaminergic neuron survival was assayed 24 hours later. The data are presented as the means ± SEM (biological replicates n=13; 3 independent experiments). \*\*\**p* < 0.001 vs NSC and F^PD^ (One-way ANOVA followed by Tukey’s test). (**C**) Low- and high-magnification (dashed brown squares) images of DIV7 midbrain cultures exposed for 24 hours to MCM from F and TPF^PD^-stimulated or unstimulated cells (NSC). Brown arrows point to dystrophic cell bodies and processes with varicosities. Scale Bar = 100 µm. (**D**) Survival of TH+ neurons in DIV7 midbrain cultures exposed for 24 hours to MCM from F^PD^ and TPF^PD^-stimulated or unstimulated cells (NSC). In sister cultures, neurons were exposed to MCM from F^PD^ and TPF^PD^-stimulated microglial cells treated with the xCT inhibitor Sulfasalazine (250 µM) during the 24 hours washout period. In additional sister cultures, neurons exposed to MCM from F^PD^ and TPF^PD^-stimulated microglial cells were cotreated with the *N*-methyl-D-aspartate (NMDA) receptor pore blocker MK-801 (2 µM). The data are presented as the means ± SEM (biological replicates n=5-8; 2 independent experiments). *** *p* < 0.001 vs. NSC or F^PD^ (One-way ANOVA followed by pairwise multiple comparisons using the Holm-Sidak method). (**E**) TH+ dopaminergic neuron survival is inversely correlated with the concentration of glutamate within the transferred MCM (linear regression analysis).

To further strengthen our hypothesis, we exposed F^PD^- and TPF^PD^-stimulated microglial cells to the xCT inhibitor Sulfasalazine (250 µM) during the 24 hour of the washout step and subsequently transferred the resulting MCM onto midbrain neurons. Sulfasalazine treatment of stimulated microglial cells completely abrogated MCM-associated dopaminergic toxicity, suggesting an essential role of xCT in the neurotoxic activity of chronic-type inflammatory microglial cells (Fig. 7D). Finally, to establish that TPF^PD^-activated microglial cells induce dopaminergic cell death through glutamate-dependent excitotoxic mechanisms, we treated MCM-exposed midbrain cultures with MK-801 (2 µM), an inhibitor of the glutamate receptor *N*-methyl-D-aspartate receptor (NMDAR). As shown in Figure 7D, MK-801 treatment fully protected against MCM-associated damage, indicating that chronic-type inflammatory microglial cells promote dopaminergic neuronal death by excitotoxicity.

## Discussion

In this study, we investigated the inflammatory properties of PD patient-derived αSYN fibrils on microglial cells and explored their capacity to induce a unique immune response when combined with TNFα and prostaglandin E_2_ (PGE_2_), two inflammatory cues commonly elevated in chronic inflammatory diseases and in PD patients. We found that fibrils derived from the PMCA of PD patient tissue have stronger inflammatory potency than recombinant human αSYN fibrils generated *de novo* in cultured mouse microglia suggesting a relationship between the structure of aggregated αSYN and the macrophage immune response. Importantly, the inflammatory response elicited by PD patient-derived αSYN fibrils on microglia was profoundly modulated in the presence of TNFα and PGE_2_ with lower cytokine but higher glutamate release responsiveness. This “chronic-type” inflammatory stimulation results in specific molecular reprogramming of microglial cells that depart from classically activated and proinflammatory M1 microglia and is characterized by seemingly increased iron retention and excitotoxic capacities.

Microglia, the most abundant resident macrophages of the central nervous system (CNS), play important roles in the maintenance of tissue homeostasis [64-66]. However, upon neuronal injury, microglia are rapidly activated and mount an inflammatory response that, if persistent, can prove neurotoxic [67-69]. In PD, protracted microglial cell activation is a neuropathological characteristic that accompanies neurodegeneration and deposition of pathological αSYN [70]. Protein aggregates are common features of several age-related neurodegenerative diseases and have been linked to microglial cell activation and the ensuing inflammatory response through ligation of pattern recognition receptors (PRRs) [5]. Mounting evidence suggests that αSYN aggregates, which are eventually released from neurons as part of the prion-like propagation mechanism of the synucleinopathies across neuronal networks, are a major pathological trigger of microglial activation and inflammation in PD [52, 71]. Yet, the importance of the respective PRRs in this process is still a subject of debate. While some authors have noted a key role for TLR2, others have found that TLR4 may also be important [53, 72-74]. The discrepancy between these results is likely to stem from the type and purity of the αSYN assemblies used in different studies. Recently, using distinct well-characterized recombinant human αSYN assemblies generated *de novo*, we found that TLR2 and TLR5, but not TLR4, were indispensable for monomeric and oligomeric αSYN to efficiently activate the NOD-like receptor pyrin domain containing 3 (NLRP3) inflammasome and IL1β release from microglial cells [8]. Equal concentrations of αSYN fibrillar polymorphs were unable to evoke IL1β release during the same stimulation period [8]. Thus, these data strongly suggest that the structural characteristics of αSYN assemblies are major determinants of their innate immunological properties. In support of this view, we showed here that fibrils derived from PD patients (F^PD^) elicited a greater inflammatory response than fibrils assembled *de novo* but a lower response than fibrils amplified from DLB patients (F^DLB^). Strikingly, whereas F^S^ (2 µM) was unable to induce IL1β release in unprimed microglial cells [8], we found that lower concentrations of F^PD^ (1.5 µM) induced a robust IL1β response (Fig. 3B). Since F^PD^ are structurally different from F and F [75, 76], our data further support the view that αSYN assemblies with different structures have distinct inflammatory potency on microglial cells. The structure-function relationship depicted here for the microglia-associated immune response is reminiscent of that documented for neuronal deposition of pathological αSYN and toxicity [15, 77]. Indeed, we recently reported that structurally distinct αSYN fibrils assembled *de novo* or derived from PD, DLB and MSA patients result in different synucleinopathy-associated characteristics and toxicity in cultured cells and after inoculation in rodent brains [12, 14, 15]. The different biological properties between patient-derived and *de novo* assembled fibrils may originate from the different surfaces that distinct fibrillar polymorphs expose to the solvent which define their differential binding to cell surface proteins, seeding propensity in cells and turnover [12, 16]. Hence, it is likely that the capacity of the F polymorph to evoke a stronger inflammatory response than its recombinant counterpart might be linked to its ability to bind with higher affinity to innate immune receptors or to activate different types of receptors. However, the binding capacity of F to different TLRs or other innate immune receptors has yet to be established.

A major contribution of our study is the demonstration that the F^PD^-induced microglial inflammatory response is profoundly altered when associated with the proinflammatory cytokine TNFα and PGE_2_, an arachidonic acid-derived lipid modulator of inflammation. As part of this study, we further showed that under such combinatory stimulation (so called TPF^PD^), microglial cells engage a distinct molecular program from that of LPS-induced proinflammatory M1 cells. TNFα and PGE_2_, are common immune-related factors featuring chronic inflammatory conditions [78, 79]. In PD, a neurodegenerative disease associated with chronic neuroinflammatory processes, the expression level of TNFα and the PGE_2_-producing enzyme cyclooxygenase 2 (COX2) were shown to increase [80-82]. Moreover, compelling experimental evidence supports an independent pathogenic role of these inflammatory factors, suggesting that in addition to being part of a large panel of effector molecules derived from activated glial cells or injured neurons, these inflammatory cues are instrumental in neuronal cell death [83, 84]. Hence, although αSYN aggregates are considered primary and master disease-associated inflammatory triggers in PD, other immune-related factors, as part of a global network of neuroinflammatory processes [4], may significantly contribute to regulating microglia-associated inflammation and toxicity. Indeed, it is now well established that microglial cells, and tissue macrophages in general, are very plastic and can adopt a wide variety of activation phenotypes associated with specific functions. Analysis and network modeling of transcriptional regulation during human macrophage activation elicited by a diverse set of stimuli have clearly established a spectrum model of macrophage activation that reassesses and extends the prevailing M1 (proinflammatory or classically activated) versus M2 (anti-inflammatory or alternatively activated) polarization model [21]. Notably, macrophages exposed to the combination of chronic inflammation-associated factor TNFα, PGE_2_, and TLR2 ligands (the so-called “TPP” stimulation) feature specific molecular modules not found in IFNγ-(M1) or IL4-(M2) stimulated cells and consequently express specific cell surface markers and cytokines. However, the findings obtained in human macrophages might not be generalizable to murine microglia. Indeed, we found that the regulation of TPP-specific markers (CD25, CXCL5 and IL1α) in mouse microglial cells was markedly different from that in human macrophages highlighting important macrophage identity- and/or species-related specificity, as previously reviewed [85]. Yet, despite these differences, our results clearly show that, as documented in human macrophages, chronic-type inflammatory stimulation of microglia is associated with specific transcriptional reprogramming departing from classically activated M1 cells. Interestingly, some of the molecular pathways characterizing chronic-type inflammatory stimulation were consistent with previously defined functional properties. In particular, the finding that TPF^PD^- stimulated cells feature higher xCT-encoding *Slc7a11* gene expression is consistent with the enhanced responsiveness of these cells to glutamate release. Importantly, a similar upregulation of *Slc7a11* was detected in human microglial-like cells (iMGs) exposed to TPF^PD^ arguing that this chronic-type inflammatory-associated regulation is conserved across mouse and human species. While releasing glutamate, the xCT antiporter internalizes cystine, which is needed for glutathione synthesis. In line with this scenario, we further observed that TPF^PD^-polarized microglial cells are characterized by the specific upregulation of several genes (*Gss, Gclc* and *Gclm*) encoding rate-limiting enzymes involved in glutathione synthesis. The cystine-glutamate antiporter xCT has been previously shown to contribute to the proinflammatory phenotype of primary microglial cells and to be involved in amyotrophic lateral sclerosis (ALS)-and Alzheimer’s disease-related pathogenesis [86, 87]. In particular, decreased production of specific microglial proinflammatory/neurotoxic factors including nitric oxide, TNFα and IL6 was observed in xCT-deleted microglial cells exposed to LPS [86]. Our data showing decreased TNFα and IL6 production concomitant with increased *Slc7a11* gene expression and glutamate release in TPF^PD^-stimulated microglia suggest that the regulatory mechanisms of proinflammatory cytokine expression induced by xCT differ between different microglial activation states. Hence, the importance of our finding in the context of PD-related neuroinflammation stems from the fact that glutamate has the potential to generate low-level excitotoxic insults [88] that may contribute to neuronal cell death [89]. In line with this, we provide compelling evidence that xCT-dependent glutamate release from chronic-type inflammatory microglia results in NMDAR overactivation-mediated excitotoxic dopaminergic cell death. In the field of PD, previous studies have reported an increase in xCT expression upon injection of 6-hydroxydopamine (6-OHDA) in hemi-Parkinson rats [90]. Moreover, neuroprotection has been documented in xCT-null mice injected with 6-OHDA [91]. Yet, inhibition of xCT by genetic targeting does not protect mice from MPTP-induced dopaminergic cell loss [92]. Thus, although debated, the role of xCT in PD pathogenesis is still unclear. Notably, its pathogenic role has never been investigated in PD animal models featuring degenerative synucleinopathy.

Apart from glutathione metabolism, the identification of the TPF^PD^-specific ferroptotic pathway in our KEGG analysis was surprising given the fact that we did not observe microglial cell death upon exposure to TPF^PD^. It has been documented that macrophages display significant defiance against ferroptotic stimulation and death, particularly when polarized into classically activated M1 state [93]. Yet, as part of the set of genes involved in this pathway, TPF stimulation was associated with elevated expression of iron-related transport proteins coupled with increased intracellular iron-storage capacity. Increased iron retention is a well-known function of inflammatory macrophages that serves as a primary defense mechanism against pathogens through extracellular iron starvation. Yet, macrophage iron uptake fulfills additional important role as it is an essential metal that fine-tunes metabolic changes associated with immune activation thereby contributing directly to macrophage function and plasticity [94]. A previous report indicated that M1-type microglia preferentially acquire iron from non-Transferrin-bound iron (NTBI), whereas M2 polarized cells promote TBI uptake [95]. Our data are consistent with this general scheme since LPS-induced microglial activation was associated with concomitant increases and decreases in the gene expression of *Slc11a2* (encoding divalent metal transporter 1; DMT1) and *Tfrc* (encoding the Transferrin receptor), respectively. Remarkably, however, TPF^PD^-stimulated cells were characterized by increased gene expression of both *Slc11a2* and *Tfrc*, indicating a unique iron transport phenotype featuring both M1 and M2 properties. The presumably greater iron retention capacity of TPF^PD^- stimulated cells than of LPS-polarized microglia is further supported by the specific upregulation of the gene encoding for ZIP14 (*Slc39a14*), a complex broad-scope metal-ion transporter of zinc and non-transferrin-bound iron. Coupled with increased iron uptake capacity, TPF^PD^-polarized microglia further displayed a high decrease in *Slc40a1* gene expression (encoding the iron extruder Ferroportin) as well as a robust increase in *Fth1*, which encodes the iron storage Ferritin heavy chain. As for *Slc7a11* gene regulation, TPF^PD^-elicited *Tfrc* and *Slc11a2* upregulated gene expression was conserved in human iMGs confirming the relevance of our findings in humans. Taken together, these data suggest that microglial cells polarized under TPF^PD^-associated chronic-type inflammation feature both M1- and M2-associated iron metabolism regulation and propose that they might be particularly iron retentive compared to M1 and M2 inflammatory cells. Relevant to our findings, is the observation that iron accumulates prematurely and preferentially in microglial cells rather than dopaminergic neurons in a nonhuman primate model of degenerative synucleinopathy induced by intranasal administration of human αSYN preformed fibrils (PFFs) [96]. Interestingly, recent data obtained from a model of human induced pluripotent stem cell-derived microglia grown in a tri-culture system with neurons and astrocytes have shown that iron-loaded microglial cells not only are susceptible to ferroptotic cell death but also prone to inducing neuronal toxicity through a mechanism that has yet to be determined [95]. Strikingly, single-cell transcriptomic and KEGG pathway analyses in this model system demonstrated that ferroptosis and glutathione metabolism were among the pathways most affected in microglial cells exposed to ferroptotic conditions [97]. These features are reminiscent of our data obtained in TPF^PD^-stimulated primary microglial cells suggesting that chronic-type inflammatory stimulation results in a microglial phenotypic shift toward a strong iron-retentive state, which may eventually favor their ferroptotic death and neurotoxicity. Alternatively, microglial iron loading could induce oxidative stress while decreasing their phagocytic activity and release capacity of inflammatory mediators [98]. Altogether these data suggest that chronic-type inflammatory microglia hold a particularly aggressive phenotype characterized by both increased toxicity and loss of protective/healing properties.

It is now well established that in response to environmental cues, such as inflammatory stimuli, microglia shift their metabolism as part of an adaptative mechanism that drives specific effector functions. Notably, TLR-related proinflammatory stimulation (e.g., LPS) induces a metabolic shift from oxidative phosphorylation to glycolysis, a phenomenon known as the Warburg effect initially characterized in tumor cells [59, 99]. Consistent with this, we found that LPS stimulation of microglia resulted in a decreased α-ketoglutarate/succinate and fumarate/succinate ratios, which are indicative of tricarboxylic acid (TCA) cycle breaks. Importantly, the immune response evoked by TPF stimulation was associated with similar metabolic changes suggesting that from a respiratory point of view this chronic-type inflammation shares common features with classical M1-polarized cells. Surprisingly, even though TPF^PD^-stimulated cells release more glutamate than LPS-exposed cells, we observed similar intracellular levels of glutamate under these two inflammatory conditions. Therefore, we posit that the greater capacity of TPF^PD^-polarized cells to extrude glutamate is likely linked to increased expression of xCT rather than enhanced glutamate synthesis pathway activity.

In summary, we documented that PD patient-derived αSYN fibrils hold robust inflammatory properties on microglial cells. Furthermore, we showed that when combined with TNFα and PGE_2_, this disease-associated αSYN polymorph polarizes microglia into a specific “chronic-type” inflammatory activation state and phenotype featuring both M1- and M2-related traits and exhibiting overwhelming neurotoxic activities. Of note, composed M1/M2 polarization assignment has been previously documented in animal models of traumatic brain injury, ALS and glioma-associated neurological disease suggesting that our model system may, in part, recapitulate more closely some of the complex innate inflammatory mechanisms *in vivo* [20, 100-102]. In support of this view, we identified few overlaps between TPF^PD^-stimulated microglia *in vitro* and the recently characterized disease-associated microglia (DAM) isolated from mouse models of Alzheimer’s disease and ALS [103]. In particular, few of the genes assigned to DAMs, including B2m, Lyz2 and Cst7, were found to be differentially expressed in TPF^PD^-polarized microglia (data not shown). Nonetheless, the global gene expression overlaps between the DAM transcriptome and that of LPS- or TPF^PD^-derived cells was modest (53 and 50 genes, respectively), highlighting significant differences in genomic expression patterns between *in vitro* and *in vivo* microglia cells, as previously reported [104]. Yet, although we are well aware that the model described here is not a phenocopy of activated microglial cells in the brains of PD patients, it may serve as a general framework for exploring and understanding the disease-associated mechanisms underlying complex inflammatory-induced signal integration that shape microglial cell activation and function. Hence, such knowledge should facilitate the identification of new potential therapeutic targets and may lead to the discovery of biomarkers.

## Ethics approval and consent to participate

The protocol used for collecting human peripheral blood mononuclear cells was approved by an ethical committee affiliated with French regulation authorities (CCPPRB Groupe Hospitalier Pitié-Salpêtrière, Paris, France). Blood samples were collected from a donor who provided signed informed consent and who was included in an INSERM-sponsored clinical study. Experimental procedures in mice were authorized by the Committee on the Ethics of Animal Experiments Charles Darwin N°5 under the registration number Ce5/2017/005.

## Consent for publication

Not applicable.

## Availability of data and materials

The datasets supporting the conclusions of this article are included within the article and its additional file.

## Authors’ contributions

C.Y. and S.H. designed all the experiments; C.Y., L.G., J.S.-D. and P.B. performed the experiments and analyzed the data with the assistance of P.P.M. and S.H.; A.F. and R.M. generated and characterized all the α-Synuclein assemblies and provided advice; and C.Y. and S.H. wrote the manuscript with input from all the coauthors.

## Abbreviations

ACM: astrocyte-conditioned medium
aSYN: alpha-Synuclein
DIV: day *in vitro*
DLB: dementia with Lewy bodies
DN: dopaminergic neuron
DAM: disease-associated microglia
DMEM^S^: supplemented Dulbecco’s modified Eagle medium
F^S^: *de novo*-generated aSYN fibrils
F^PD^: PD patient-derived aSYN fibrils
F^DLB^: DLB patient-derived aSYN fibrils
iMG: induced microglia-like cell
L-15: Leibovitz’s L-15 medium
LP: Lewy pathology
LPS: lipopolysaccharide
M1: type-1 macrophage
M2: type-2 macrophage
MCM: microglia-conditioned medium
MSA: multiple system atrophy
Nb^S^: supplemented neurobasal-A medium
ND: nondifferentiated
NSC: nonstimulated cells
PD: Parkinson’s disease
PGE_2_: prostaglandin E_2_
PMCA: protein misfolding cyclic amplification
ROS: reactive oxygen species
TEM: transmission electron microscopy TLR toll-like receptor
TNFα: tumor necrosis factor alpha
TPF^PD^: TNFα, PGE_2_, F^PD^
TPP: TNFα, PGE_2_, Pam3CSK4

## Acknowledgements

The authors would like to thank Prof. Jean-Christophe Corvol and the Clinical Investigation Center for Neurosciences of the Pitié-Salpêtrière Hospital (CIC 1421, Paris, France) for their help in collecting human blood samples. We are indebted to Prof. Steve Gentleman at the Neuropathology Unit, Division of Brain Sciences, Department of Medicine, Imperial College London for providing us with PD and DLB patient brain tissues. We also thank Dr. Luc Bousset for his invaluable advice in preparing αSyn assemblies *in vitro*. Part of this work was carried out on the CELIS cell culture, the iGenSeq and the Data and Analysis core facilities of ICM. We gratefully acknowledge Yannick Marie, Justine Guegan and David Akbar for helpful discussions and advice on the technical aspects of the project. We would also like to thank Farid Ichou and Maha Ponnaiah from the ICANalytics core facility (ICAN Institute, Salpêtrière Hospital, Paris France) for their help in sample preparation and processing for metabolomics and bioinformatics data analysis and Tracy Bellande for expert technical assistance. Finally, we thank the patients and caregivers for their participation and contribution to this study.

## References

1. Michel PP, Hirsch EC, Hunot S. Understanding Dopaminergic Cell Death Pathways in Parkinson Disease. Neuron. 2016; 90(4):675–691.

2. Brundin P, Melki R, Kopito R. Prion-like transmission of protein aggregates in neurodegenerative diseases. Nat Rev Mol Cell Biol. 2010; 11(4):301–7.

3. Dehay B, Bourdenx M, Gorry P, Przedborski S, Vila M, Hunot S, et al., Targeting α-synuclein for treatment of Parkinson’s disease: Mechanistic and therapeutic considerations. Lancet Neurol. 2015; 14(8):855–866.

4. Hirsch EC, Hunot S. Neuroinflammation in Parkinson’s disease: A target for neuroprotection? Lancet Neurol. 2009; 8(4):382–397.

5. Heneka M T, Carson MJ, Khoury JE, Landreth GE, Brosseron F, Feinstein DL, et al., Neuroinflammation in Alzheimer’s disease. Lancet Neurol. 2015; 14(4):388–405.

6. Lim S, Chun Y, Lee JS, Lee S-J. Neuroinflammation in Synucleinopathies. Brain Pathol. 2016; 26(3):404–409.

7. Panicker N, Sarkar S, Harischandra DS, Neal M, Kam T-I, Jin H, et al., Fyn kinase regulates misfolded α-synuclein uptake and NLRP3 inflammasome activation in microglia. J Exp Med. 2019; 216(6):1411–1430.

8. Scheiblich H, Bousset L, Schwartz S, Griep A, Latz E, Melki R, et al., Microglial NLRP3 Inflammasome Activation upon TLR2 and TLR5 Ligation by Distinct α-Synuclein Assemblies. J Immunol. 2021; 207(8):2143–2154.

9. Scheiblich H, Dansokho C, Mercan D, Schmidt SV, Bousset L, Wischhof L, et al., Microglia jointly degrade fibrillar alpha-synuclein cargo by distribution through tunneling nanotubes. Cell. 2021; 184(20):5089–5106.

10. Bousset L, Pieri L, Ruiz-Arlandis G, Gath J, Jensen PH, Habenstein, et al., Structural and functional characterization of two alpha-synuclein strains. Nat Commun. 2013; 4(1):2575.

11. Gribaudo S, Tixador P, Bousset L, Fenyi A, Lino P, Melki R, et al., Propagation of α-Synuclein strains within human reconstructed neuronal network. Stem Cell Rep. 2019; 12(2):230–244.

12. Shrivastava AN, Bousset L, Renner M, Redeker V, Savistchenko J, Triller A, et al., Differential membrane binding and seeding of distinct α-Synuclein fibrillar polymorphs. Biophys J. 2020; 118(6):1301–1320.

13. Peelaerts W, Bousset L, Van der Perren A, Moskalyuk A, Pulizzi R, Giugliano M, et al., α-Synuclein strains cause distinct synucleinopathies after local and systemic administration. Nature. 2015; 522(7556):340–344.

14. Rey NL, Bousset L, George S, Madaj Z, Meyerdirk L, Schulz E, et al., α-Synuclein conformational strains spread, seed and target neuronal cells differentially after injection into the olfactory bulb. Acta Neuropathol Commun. 2019; 7(1):221.

15. Van der Perren A, Gelders G, Fenyi A, Bousset L, Brito F, Peelaerts W, et al., The structural differences between patient-derived α-synuclein strains dictate characteristics of Parkinson’s disease, multiple system atrophy and dementia with Lewy bodies. Acta Neuropathol. 2020; 139(6):977–1000.

16. Landureau M, Redeker V, Bellande T, Eyquem S, Melki R. The differential solvent exposure of N-terminal residues provides “fingerprints” of alpha-synuclein fibrillar polymorphs. J Biol Chem. 2021; 296:100737.

17. Song S-Y, Kim I-S, Koppula S, Park J-Y, Kim B-W, Yoon S-H, et al., 2-Hydroxy-4-Methylbenzoic Anhydride Inhibits Neuroinflammation in Cellular and Experimental Animal Models of Parkinson’s Disease. Int J Mol Sci. 2020; 21(21):8195.

18. Tu D, Gao Y, Yang R, Guan T, Hong J-S, Gao H-M. The pentose phosphate pathway regulates chronic neuroinflammation and dopaminergic neurodegeneration. J Neuroinflammation. 2019; 16(1):1– 17.

19. Perry VH, Nicoll JAR, Holmes C. Microglia in neurodegenerative disease. Nat Rev Neurol. 2010; 6(4):193–201.

20. Chiu IM, Morimoto ETA, Goodarzi H, Liao JT, O’Keeffe S, Phatnani HP, et al., A Neurodegeneration-Specific Gene-Expression Signature of Acutely Isolated Microglia from an Amyotrophic Lateral Sclerosis Mouse Model. Cell Rep. 2013; 4(2):385–401.

21. Xue J, Schmidt SV, Sander J, Draffehn A, Krebs W, Quester I, et al., Transcriptome-Based Network Analysis Reveals a Spectrum Model of Human Macrophage Activation. Immunity. 2014; 40(2):274– 288.

22. Ciofani M, Madar A, Galan C, Sellars M, Mace K, Pauli F, et al., A Validated Regulatory Network for Th17 Cell Specification. Cell. 2012; 151(2):289–303.

23. Marino MW, Dunn A, Grail D, Inglese M, Noguchi Y, Richards E, et al., Characterization of tumor necrosis factor-deficient mice. Proc Natl Acad Sci USA. 1997; 94(15):8093–8098.

24. Popov A, Abdullah Z, Wickenhauser C, Saric T, Driesen J, Hanisch F-G, et al., Indoleamine 2,3- dioxygenase–expressing dendritic cells form suppurative granulomas following Listeria monocytogenes infection. J Clin Invest. 2006; 116(12):3160–3170.

25. Reiling N, Hölscher C, Fehrenbach A, Kröger S, Kirschning CJ, Goyert S, et al., Cutting Edge: Toll-Like Receptor (TLR)2- and TLR4-Mediated Pathogen Recognition in Resistance to Airborne Infection with *Mycobacterium tuberculosis*. J Immunol. 2002; 169(7):3480–3484.

26. Shay JES, Celeste Simon M. Hypoxia-inducible factors: Crosstalk between inflammation and metabolism. Sem Cell Dev Biol. 2012; 23(4):389–394.

27. Doorn KJ, Moors T, Drukarch B, van de Berg WD, Lucassen PJ, van Dam, A-M. Microglial phenotypes and toll-like receptor 2 in the substantia nigra and hippocampus of incidental Lewy body disease cases and Parkinson’s disease patients. Acta Neuropathol Commun. 2014; 2(1):90.

28. Hunot S, Boissière F, Faucheux B, Brugg B, Mouatt-Prigent A, Agid Y et al., Nitric oxide synthase and neuronal vulnerability in parkinson’s disease. Neuroscience. 1996; 72(2):355–363.

29. Wu Y-R, Feng I-H, Lyu R-K, Chang K-H, Lin Y-Y, Chan H et al., Tumor necrosis factor-α promoter polymorphism is associated with the risk of Parkinson’s disease. Am J Med Genet. 2007; 144B(3):300–304.

30. Sepulveda-Diaz JE, Ouidja MO, Socias SB, Hamadat S, Guerreiro S, Raisman-Vozari R, et al., A simplified approach for efficient isolation of functional microglial cells: Application for modeling neuroinflammatory responses *in vitro*: Simplified Approach for Microglia Isolation. Glia. 2016; 64(11):1912–1924.

31. Tourville A, Akbar D, Corti O, Prehn JHM, Melki R, Hunot S, et al., Modelling α-Synuclein Aggregation and Neurodegeneration with Fibril Seeds in Primary Cultures of Mouse Dopaminergic Neurons. Cells. 2022; 11(10):1640.

32. Pozzi D, Ban J, Iseppon F, Torre V. An improved method for growing neurons: Comparison with standard protocols. J Neurosci Methods. 2017; 15:280:1–10.

33. Rook GAW, Steele J, Umar S, Dockrell HM. A simple method for the solubilisation of reduced NBT, and its use as a colorimetric assay for activation of human macrophages by γ-interferon. J Immunol Methods. 1985; 82(1):161–167.

34. Love MI, Huber W, Anders S. Moderated estimation of fold change and dispersion for RNA-seq data with DESeq2. Genome Biol. 2014; 15(12):550.

35. Chen EY, Tan CM, Kou Y, Duan Q, Wang Z, Meirelles GV, et al., Enrichr: Interactive and collaborative HTML5 gene list enrichment analysis tool. BMC Bioinform. 2013; 14(1):128.

36. Kuleshov MV, Jones MR, Rouillard AD, Fernandez NF, Duan Q, Wang Z, et al., Enrichr: A comprehensive gene set enrichment analysis web server 2016 update. Nucleic Acids Res. 2016; 44(W1):W90–W97.

37. Xie Z, Bailey A, Kuleshov MV, Clarke DJB., Evangelista JE, Jenkins SL, et al., Gene Set Knowledge Discovery with Enrichr. Curr Protoc. 2021; 1(3).

38. Garali I, Adanyeguh IM, Ichou F, Perlbarg V, Seyer A, Colsch B, et al., A strategy for multimodal data integration: Application to biomarkers identification in spinocerebellar ataxia. Brief Bioinform. 2018; 19(6):1356–1369.

39. Giacomoni F, Le Corguille G, Monsoor M, Landi M, Pericard P, Petera M, et al., Workflow4Metabolomics: A collaborative research infrastructure for computational metabolomics. Bioinformatics. 2015; 31(9):1493–1495.

40. Smith CA, Want EJ, O’Maille G, Abagyan R, Siuzdak G. XCMS: Processing Mass Spectrometry Data for Metabolite Profiling Using Nonlinear Peak Alignment, Matching, and Identification. Anal Chem. 2006; 78(3):779–787.

41. Tautenhahn R, Patti GJ, Rinehart D, Siuzdak G. XCMS Online: A Web-Based Platform to Process Untargeted Metabolomic Data. Anal Chem. 2012; 84(11):5035–5039.

42. Dunn WB, Broadhurst D, Begley P, Zelena E, Francis-McIntyre S, Anderson N, et al., Procedures for large-scale metabolic profiling of serum and plasma using gas chromatography and liquid chromatography coupled to mass spectrometry. Nat Protoc. 2011; 6(7):1060–1083.

43. Want EJ, Wilson ID, Gika H, Theodoridis G, Plumb RS, Shockcor J, et al., Global metabolic profiling procedures for urine using UPLC–MS. Nat Protoc. 2010; 5(6):1005–1018.

44. Dunn WB, Wilson ID, Nicholls AW, Broadhurst D. The importance of experimental design and QC samples in large-scale and MS-driven untargeted metabolomic studies of humans. Bioanalysis. 2012; 4(18): 2249–2264.

45. Veselkov KA, Vingara LK, Masson P, Robinette SL, Want E, Li JV, et al., Optimized Preprocessing of Ultra-Performance Liquid Chromatography/Mass Spectrometry Urinary Metabolic Profiles for Improved Information Recovery. Anal Chem. 2011; 83(15):5864–5872.

46. Saeed AI, Sharov V, White J, Li J, Liang W, Bhagabati N, et al., TM4: A Free, Open-Source System for Microarray Data Management and Analysis. BioTechniques. 2003; 34(2):374–378.

47. Benjamini Y, Hochberg Y. Controlling the False Discovery Rate: A Practical and Powerful Approach to Multiple Testing. J. R. Stat. Soc. 1995; 57(1):289–300.

48. Ohgidani M, Kato TA, Setoyama D, Sagata N, Hashimoto R, Shigenobu K, et al., Direct induction of ramified microglia-like cells from human monocytes: Dynamic microglial dysfunction in Nasu-Hakola disease. Sci Rep. 2015; 4(1):4957.

49. Döring C, Regen T, Gertig U, van Rossum D, Winkler A, Saiepour N, et al., A presumed antagonistic LPS identifies distinct functional organization of TLR4 in mouse microglia. Glia. 2017; 65(7):1176– 1185.

50. dos-Santos-Pereira M, Acuña L, Hamadat S, Rocca J, González-Lizárraga F, Chehín R, et al., Microglial glutamate release evoked by α-synuclein aggregates is prevented by dopamine. Glia. 2018; 66(11):2353–2365.

51. Ginhoux F, Greter M, Leboeuf M, Nandi S, See P, Gokhan S, et al., Fate Mapping Analysis Reveals That Adult Microglia Derive from Primitive Macrophages. Science. 2010; 330(6005):841–845.

52. Choi I, Zhang Y, Seegobin S P, Pruvost M, Wang Q, Purtell K, et al., Microglia clear neuron-released α-synuclein via selective autophagy and prevent neurodegeneration. Nat Commun. 2020; 11(1):1386.

53. Fellner L, Irschick R, Schanda K, Reindl M, Klimaschewski L, Poewe W, et al., Toll-like receptor 4 is required for α-synuclein dependent activation of microglia and astroglia. Glia. 2013; 61(3):349– 360.

54. Gustot A, Gallea JI, Sarroukh R, Celej MS, Ruysschaert J-M, Raussens V. Amyloid fibrils are the molecular trigger of inflammation in Parkinson’s disease. Biochem J. 2015; 471(3):323–333.

55. Lim S, Chun Y, Lee JS, Lee SJ. Neuroinflammation in Synucleinopathies. Brain Pathol. 2016; 26(3):404–9.

56. Drier Y, Sheffer M, Domany E. Pathway-based personalized analysis of cancer. Proc Natl Acad Sci. USA. 2013; 110(16):6388–6393.

57. Lewerenz J, Hewett SJ, Huang Y, Lambros M, Gout PW, Kalivas PW, et al., The Cystine/Glutamate Antiporter System xc-in Health and Disease: From Molecular Mechanisms to Novel Therapeutic Opportunities. Antioxid Redox Signal. 2013; 18(5):522–555.

58. Rodríguez-Prados JC, Través PG, Cuenca J, Rico D, Aragonés J, Martín-Sanz P, et al., Substrate Fate in Activated Macrophages: A Comparison between Innate, Classic, and Alternative Activation. J Immunol. 2010; 185(1):605–614.

59. Orihuela R, McPherson CA, Harry GJ. Microglial M1/M2 polarization and metabolic states: Microglia bioenergetics with acute polarization. Br J Pharmacol. 2016; 173(4):649–665.

60. Angajala A, Lim S, Phillips JB, Kim J-H, Yates C, You Z, et al., Diverse roles of mitochondria in immune responses: novel insights into immuno-metabolism. Front Immunol. 2018; 9:1605.

61. Galatro TF, Holtman IR, Lerario AM, Vainchtein ID, Brouwer N, Sola PR, et al., Transcriptomic analysis of purified human cortical microglia reveals age-associated changes. Nat Neurosci. 2017; 20(8):1162–1171.

62. Mizutani M, Pino PA, Saederup N, Charo IF, Ransohoff RM, Cardona AE. The Fractalkine Receptor but Not CCR2 Is Present on Microglia from Embryonic Development throughout Adulthood. J Immunol. 2012; 188(1):29–36.

63. Wang J, Wang F, Mai D, Qu, S. Molecular Mechanisms of Glutamate Toxicity in Parkinson’s Disease. Front Neurosci. 2020; 14:585584.

64. Colonna M, Butovsky O. Microglia Function in the Central Nervous System During Health and Neurodegeneration. Annu Rev Immunol. 2017; 35(1):441–468.

65. Prinz M, Jung S, Priller J. Microglia Biology: One Century of Evolving Concepts. Cell. 2019; 179(2): 292–311.

66. Ransohoff RM, Perry VH. Microglial Physiology: Unique Stimuli, Specialized Responses. Ann Rev Immunol. 2009; 27(1):119–145.

67. Bodea L-G, Wang Y, Linnartz-Gerlach B, Kopatz J, Sinkkonen L, Musgrove R, et al., Neurodegeneration by Activation of the Microglial Complement-Phagosome Pathway. J Neurosci. 2014; 34(25):8546–8556.

68. Donat CK, Scott G, Gentleman SM, Sastre M. Microglial Activation in Traumatic Brain Injury. Frontiers in Aging Neurosci. 2017; 9:208.

69. Lull ME, Block ML. Microglial activation and chronic neurodegeneration. Neurotherapeutics. 2010; 7(4):354–365.

70. Ouchi Y, Yagi S, Yokokura M, Sakamoto M. Neuroinflammation in the living brain of Parkinson’s disease. Parkinsonism Relat Disord. 2009; 15:S200–S204.

71. Daniele SG, Béraud D, Davenport C, Cheng K, Yin H, Maguire-Zeiss KA. Activation of MyD88-dependent TLR1/2 signaling by misfolded α-synuclein, a protein linked to neurodegenerative disorders. Sci Signal. 2015; 8(376).

72. Codolo G, Plotegher N, Pozzobon T, Brucale M, Tessari I, Bubacco L, et al., Triggering of Inflammasome by Aggregated α–Synuclein, an Inflammatory Response in Synucleinopathies. PLoS ONE. 2013; 8(1):e55375.

73. Feng Y, Zheng C, Zhang Y, Xing C, Cai W, Li R et al., Triptolide Inhibits Preformed Fibril-Induced Microglial Activation by Targeting the MicroRNA155-5p/SHIP1 Pathway. Oxid Med Cell Longev. 2019; 1–13.

74. Kim C, Ho D-H, Suk J-E, You S, Michael S, Kang J, et al., Neuron-released oligomeric α-synuclein is an endogenous agonist of TLR2 for paracrine activation of microglia. Nat Commun. 2013; 4(1):1562.

75. Guerrero-Ferreira R, Taylor NM, Arteni AA, Kumari P, Mona D, Ringler P, et al., Two new polymorphic structures of human full-length alpha-synuclein fibrils solved by cryo-electron microscopy. Elife. 2019; 8:e48907.

76. Burger D, Fenyi A, Bousset L, Stahlberg H, Melki R. Cryo-EM structure of alpha-synuclein fibrils amplified by PMCA from PD and MSA patient brains. bioRxiv. 2021; 451588; doi: 10.1101/2021.07.08.451588.

77. Shahnawaz M, Mukherjee A, Pritzkow S, Mendez N, Rabadia P, Liu X, et al., Discriminating α- synuclein strains in Parkinson’s disease and multiple system atrophy. Nature. 2020; 578(7794):273–277.

78. Aoki T, Narumiya S. Prostaglandins and chronic inflammation. Trends Pharmacol Sci. 2012; 33(6):304–311.

79. Balkwill F. Tumor necrosis factor and cancer. Nat. Rev Cancer. 2009; 9(5):361–371.

80. Mattammal MB, Strong R, Lakshmi VM, Chung HD, Stephenson AH. Prostaglandin H Synthetase-Mediated Metabolism of Dopamine: Implication for Parkinson’s Disease. J Neurochem. 2002; 64(4):1645–1654.

81. Mogi M, Harada M, Riederer P, Narabayashi H, Fujita K, Nagatsu T. Tumor necrosis factor-α (TNF-α) increases both in the brain and in the cerebrospinal fluid from parkinsonian patients. Neurosci Lett. 1994; 165(1–2):208–210.

82. Teismann P, Tieu K, Choi D-K, Wu D-C, Naini A, Hunot S, et al., Cyclooxygenase-2 is instrumental in Parkinson’s disease neurodegeneration. Proc Natl Acad Sci USA. 2003; 100(9):5473–5478.

83. Tansey MG, Goldberg MS. Neuroinflammation in Parkinson’s disease: Its role in neuronal death and implications for therapeutic intervention. Neurobiol Dis. 2010; 37(3):510–518.

84. Teismann P, Tieu K, Cohen O, Choi D-K, Wu DC, Marks D, et al., Pathogenic role of glial cells in Parkinson’s disease. Mov Disord. 2003; 18(2):121–129.

85. Locati M, Curtale G, Mantovani A. Diversity, Mechanisms, and Significance of Macrophage Plasticity. Ann Rev Pathol. 2020; 15(1):123–147.

86. Mesci P, Zaïdi S, Lobsiger CS, Millecamps S, Escartin C, Seilhean D, et al., System xC− is a mediator of microglial function and its deletion slows symptoms in amyotrophic lateral sclerosis mice. Brain. 2015; 138(1):53–68.

87. Qin S, Colin C, Hinners I, Gervais A, Cheret C, Mallat M. System Xc- and Apolipoprotein E Expressed by Microglia Have Opposite Effects on the Neurotoxicity of Amyloid-beta Peptide 1-40. J Neurosci. 2006; 26(12):3345–3356.

88. Lavaur J, Le Nogue D, Lemaire M, Pype J, Farjot G, Hirsch EC, et al., The noble gas xenon provides protection and trophic stimulation to midbrain dopamine neurons. J Neurochem. 2017; 142(1):14–28.

89. Ambrosi G, Cerri S, Blandini F. A further update on the role of excitotoxicity in the pathogenesis of Parkinson’s disease. J Neural Transm. 2014; 121(8):849–859.

90. Massie A, Schallier A, Mertens B, Vermoesen K, Bannai S, Sato H, et al., Time-dependent changes in striatal xCT protein expression in hemi-Parkinson rats. NeuroReport. 2008; 19(16):1589–1592.

91. Massie A, Schallier A, Kim SW, Fernando R, Kobayashi S, Beck H, et al., Dopaminergic neurons of system x(c)^−^-deficient mice are highly protected against 6-hydroxydopamine-induced toxicity. FASEB J. 2011; 25(4):1359–1369.

92. Bentea E, Sconce MD, Churchill MJ, Van Liefferinge J, Sato H, Meshul CK, et al., MPTP-induced parkinsonism in mice alters striatal and nigral xCT expression but is unaffected by the genetic loss of xCT. Neurosci Lett. 2015; 593:1–6.

93. Kapralov AA, Yang Q, Dar HH, Tyurina YY, Anthonymuthu TS, Kim R, et al., Redox lipid reprogramming commands susceptibility of macrophages and microglia to ferroptotic death. Nat Chem Biol. 2020; 16(3):278–290.

94. Pereira M, Chen T-D, Buang N, Olona A, Ko J-H, Prendecki M, et al., Acute Iron Deprivation Reprograms Human Macrophage Metabolism and Reduces Inflammation In Vivo. Cell Rep. 2019; 28(2):498–511.e5.

95. McCarthy RC, Sosa JC, Gardeck AM, Baez AS, Lee C-H, Wessling-Resnick M. Inflammation-induced iron transport and metabolism by brain microglia. J Biol Chem. 2018; 293(20):7853–7863.

96. Guo J-J, Yue F, Song D-Y, Bousset L, Liang X, Tang J, et al., Intranasal administration of α-synuclein preformed fibrils triggers microglial iron deposition in the substantia nigra of Macaca fascicularis. Cell Death Dis. 2021; 12(1):81.

97. Ryan SK, Zelic M, Han Y, Teeple E, Chen L, Sadeghi M, et al., Microglia ferroptosis is regulated by SEC24B and contributes to neurodegeneration. Nat Neurosci. 2023; 26:12–26.

98. Kenkhuis B, van Eekeren M, Parfitt DA, Ariyurek Y, Banerjee P, Priller J, et al., Iron accumulation induces oxidative stress, while depressing inflammatory polarization in human iPSC-derived microglia. Stem Cell Rep. 2022; 17:1351–1365.

99. Warburg O. On respiratory impairment in cancer cells. Science. 1956; 124:269–70.

100. Kim CC, Nakamura MC, Hsieh CL. Brain trauma elicits non-canonical macrophage activation states. J Neuroinflammation. 2016; 13(1):117.

101. Morganti JM, Riparip L-K, Rosi S. Call Off the Dog(ma): M1/M2 polarization Is concurrent following traumatic brain injury. PLoS ONE. 2016; 11(1):e0148001.

102. Szulzewsky F, Pelz A, Feng X, Synowitz M, Markovic D, Langmann T, et al., Glioma-Associated Microglia/Macrophages Display an Expression Profile Different from M1 and M2 Polarization and Highly Express Gpnmb and Spp1. PLoS ONE. 2015; 10(2):e0116644.

103. Keren-Shaul H, Spinrad A, Weiner A, Matcovitch-Natan O, Dvir-Szternfeld R, Ulland TK, et al., A Unique Microglia Type Associated with Restricting Development of Alzheimer’s Disease. Cell. 2017; 169(7):1276–1290.

104. McFarland KN, Ceballos C, Rosario A, Ladd T, Moore B, Golde G, et al., Microglia show differential transcriptomic response to Aβ peptide aggregates ex vivo and in vivo. Life Sci Alliance. 2021; 4(7):e202101108.

