## Supplemental Figures S1 to S3 for "Parkinson’s disease-derived α-Synuclein assemblies combined with chronic-type inflammatory cues promote a neurotoxic microglial phenotype"

**Additional File 1.**

**Supplemantal Figures S1 to S3**

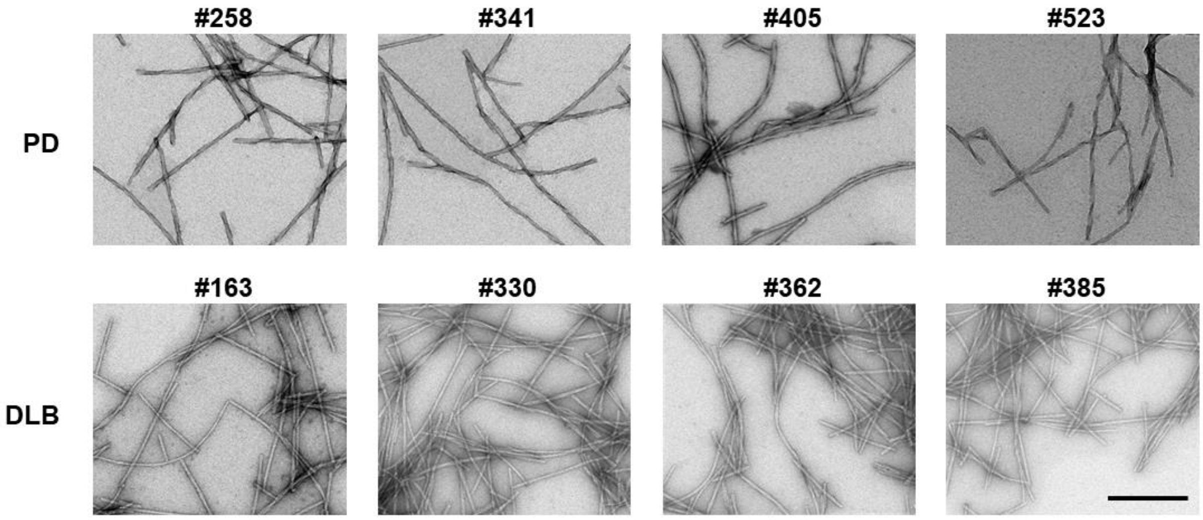


**Figure S1. Morphology of patient-derived αSYN assemblies.** Electron micrographs of αSYN assemblies generated after the 3rd cycle of amplification by PMCA from 4 PD (#258, 341, 405, 523) and 4 DLB (#163, 330, 362, 385) patients. Note the cylindrical and twisted appearance of PD-derived assemblies compared to the flatted and not-twisted shape of those derived from DLB patients. Scale bar = 200 nm.

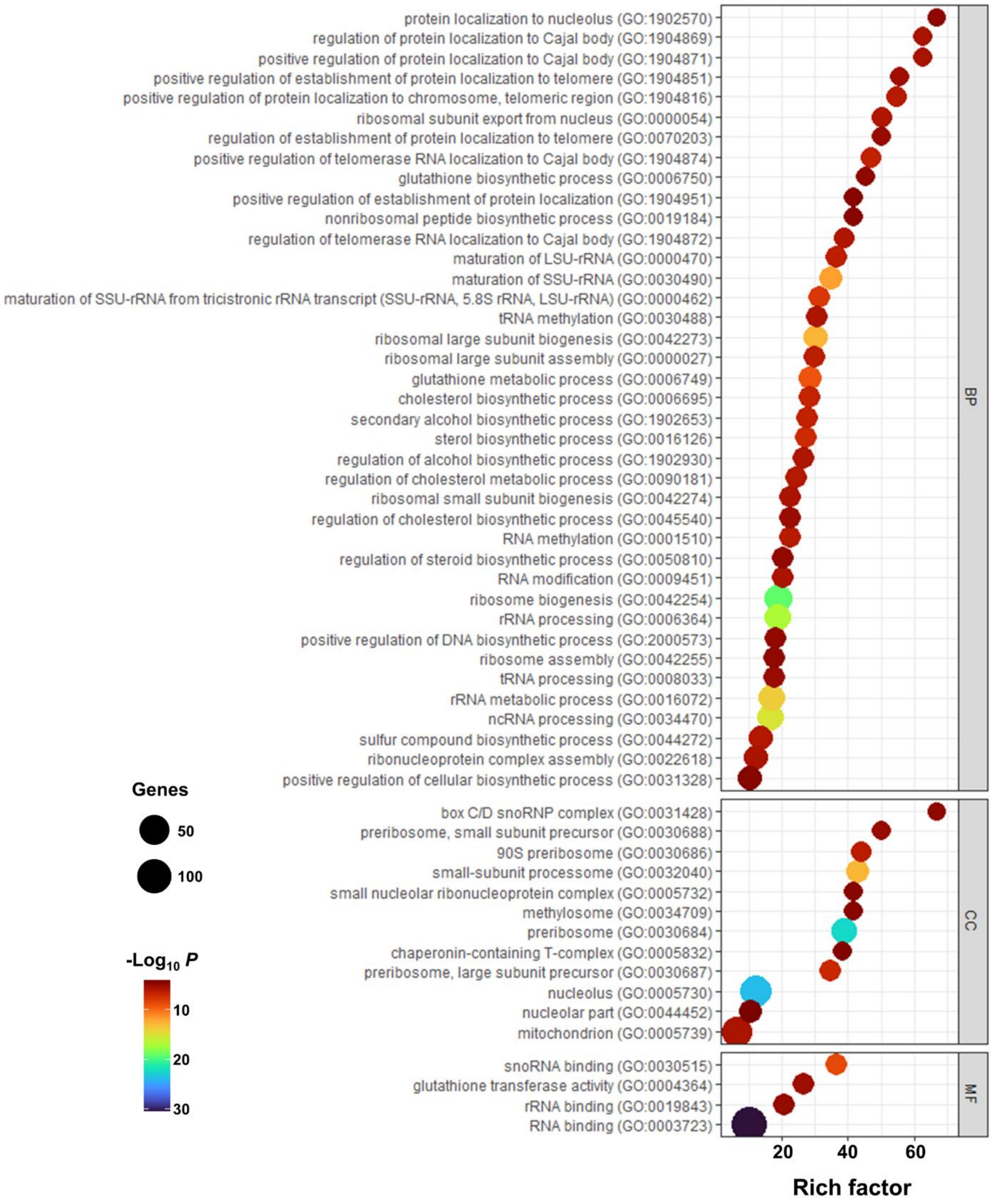


**Figure S2. TPF$^{PD}$-specific gene ontology term enrichment analysis.** Bubble chart showing the enrichment of the GO terms in DEG of TPF$^{PD}$-treated microglial cells (adjusted $p < 0.05$). Bubble size indicates the number of genes annotated in the indicated GO term enrichment. Colors represent pathway enrichment (% of overlapping genes) in TPF$^{PD}$-exposed cells.

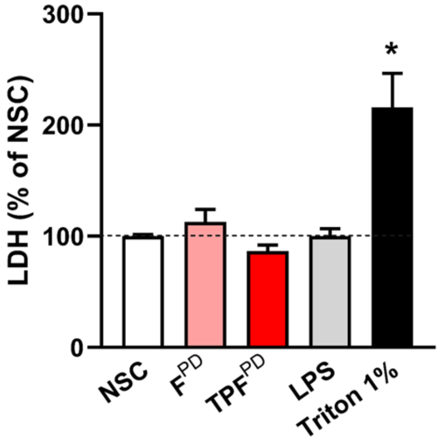


**Figure S3. TPF$^{PD}$-related chronic-type inflammatory stimulation does not induce microglial cell death.** Quantification of LDH release in microglial cells exposed or not (NSC) to F$^{PD}$ (1.5 µM), TNFα+PGE$_2$+F$^{PD}$ (TPF$^{PD}$), LPS (10 ng/mL) or 1% Triton X-100 (as a positive control). Any treatment but 1% Triton X100 result in cell membrane disruption-associated LDH release. Bars are means ± SEM (n = 3-6). *$p$ < 0.05 vs. all other conditions (Tukey's test).
